# Sustained effect of MPOA Penk neurons underlies progression through consummatory mating behavior in male mice

**DOI:** 10.64898/2026.04.03.716295

**Authors:** Yousuke Tsuneoka, Kouta Kanno, Kimiya Narikiyo, Hiromasa Funato

## Abstract

Male rodent sexual behavior progresses from appetitive to consummatory mating, implying a sustained internal motivational state. Estrogen receptor 1 (*Esr1*)-expressing neurons in the medial preoptic area (MPOA) regulate discrete mating actions, but how motivational drive is sustained across mating remains unknown. Here, we identify an MPOA *Esr1*+ neuronal subtype marked by proenkephalin (Penk) that promotes the transition from mount to intromission. Male mice exhibited a bimodal progression to consummatory mating, and *Penk*-expressing neurons represented the predominant population recruited for consummatory behaviors. In sexually active males, *Penk*+ neurons exhibited sustained Ca²⁺ dynamics. Chemogenetic manipulation selectively enhanced female-directed consummatory behaviors without affecting intermale interactions. Optogenetic activation of *Penk*+ neurons or their terminals promoted consummatory behavior over a behaviorally relevant timescale. Thus, MPOA *Penk*+ neurons provide a circuit substrate for a sustained internal state that ensures successful mating progression.

## Introduction

Male sexual behavior is a prototypal motivated behavior that unfolds in distinct phases. In rodents, an appetitive phase involving partner search and courtship typically precedes a consummatory phase characterized by mounting, intromission, and ejaculation ^1,2^. Appetitive behaviors reflect preparatory processes that establish the conditions for mating, whereas mounting, intromission, and ejaculation constitute consummatory behaviors to accomplish sexual interaction. These elements are thought to be governed by a sustained internal motivational state, or sexual arousal state, that becomes engaged during the appetitive phase and is maintained until ejaculation, thereby enabling the transition from appetitive to consummatory behavior and supporting the successful completion of the mount-intromission-ejaculation sequence ^1,3,4^. However, the neural basis of this sustained drive state that bridges appetitive actions to the execution of consummatory behaviors remains largely unknown.

The medial preoptic area (MPOA) has been recognized as a core hypothalamic hub for male sexual behavior, integrating sensory inputs, hormonal milieu, and social context ^2,5–8^. Within the MPOA, *estrogen receptor alpha* (*Esr1*)-expressing neurons are indispensable for male mating-related behaviors ^9,10^. Histological analyses and single-cell transcriptomic profiling indicate that *Esr1*-positive populations comprise multiple molecularly distinct subpopulations, raising the possibility that each subpopulation plays a specialized role across mating phases (Tsuneoka, Yoshida, et al., 2017; Moffitt et al., 2018). Consistent with this heterogeneity, activation of *Esr1/Vgat*-positive neurons elicits mounting toward females as well as males ^11,12^, and activation of dopaminergic inputs to the MPOA enhances female-directed sniffing and mounting via *Drd1*-positive cells, most of which are *Esr1*-positive ^13^. In addition, activation of *Tacr1*-positive cells, a subpopulation of *Esr1* expressing cells, can rapidly evoke mounting even toward males, toys and unreceptive females, with mounting terminating immediately upon cessation of stimulation ^14^. Thus, while neuronal populations capable of directly triggering sexual actions irrespective of an appropriate female partner have been identified, the physiologically relevant MPOA neuronal populations that can represent sustained motivational states and support the transition to and completion of consummatory mating behavior remain unknown.

Here we identify a central MPOA *Penk*-expressing neuronal subtype as a key candidate substrate for an internal state that promotes consummatory mating over behaviorally relevant timescales. In mice, mating behavior diverges into distinct trajectories that either complete copulation or stall after robust appetitive engagement. Using activity mapping, calcium dynamics, optogenetic/chemogenetic manipulation, and projection-specific manipulation, we show that Penk neurons promote the mount-to-intromission transition and activate circuits in the VTA and PAG through which distinct mating behaviors are executed.

## Results

### Bimodal progression to consummatory mating in male C57BL/6J mice

Despite unimpaired fertility in our C57BL/6J colony, we identified striking variability in male mating performance, especially the transition to the consummatory behavior. To quantify this inter-individual variation, hormonally primed female mice were introduced into the home cages of sexually naïve or experienced male mice, and mating-related behaviors were scored. The next day, copulatory plugs were detected in all paired females, confirming that all males were competent to mate. Male behavior exhibited a bimodal distribution: 13 out of 24 males ejaculated within 60 minutes, whereas the remainder did not (Figure 1a). We classified these as full mating episodes (FM) and partial mating episodes (PM), respectively. In FM, investigatory sniffing was followed by mounting, and proceeded to intromission and ejaculation (Figure 1b–1d, S1a). After ejaculation, FM males showed a marked reduction in female-directed interactions (Figure S1c).

**Figure 1.**
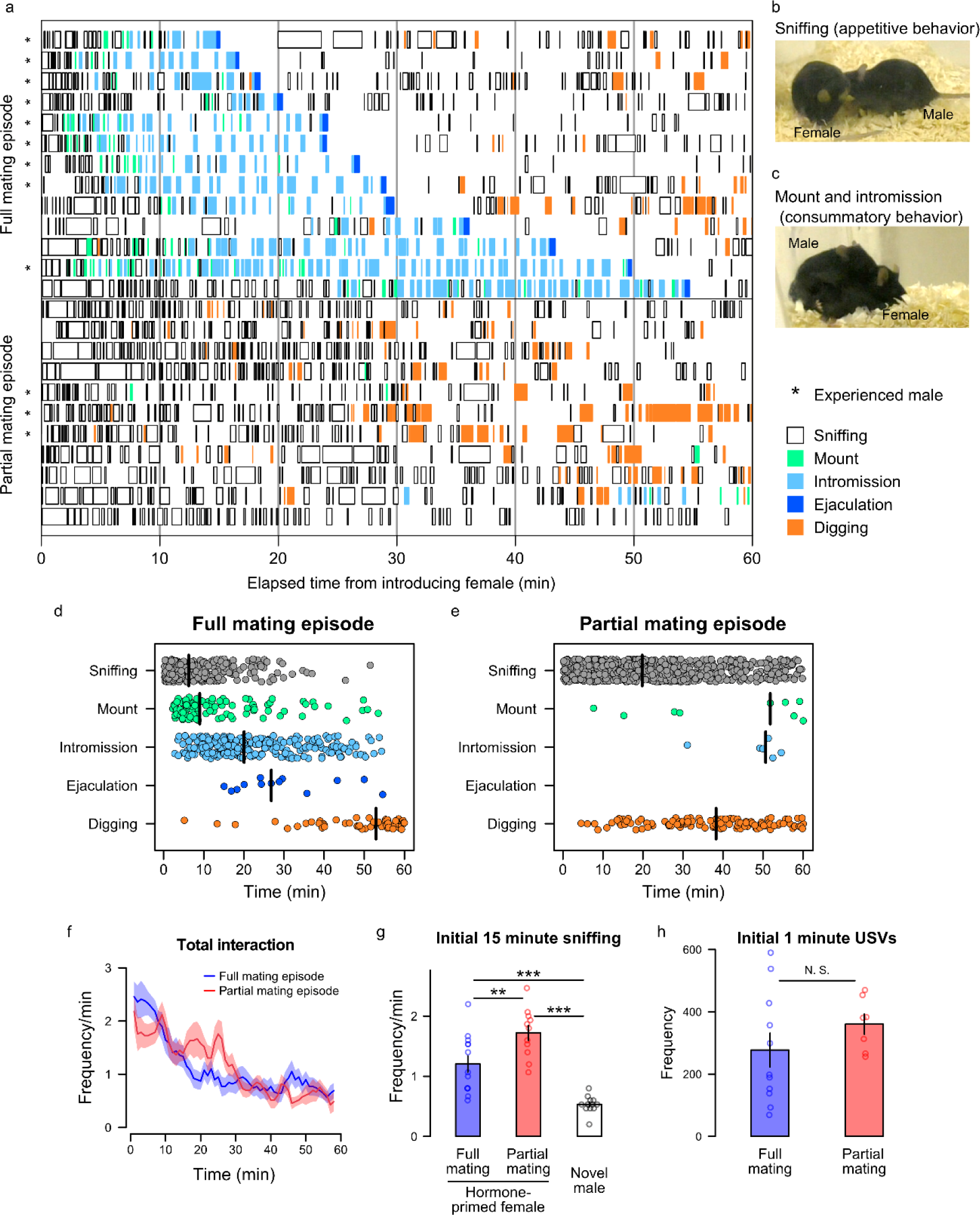
Behavioral characterization of full and partial mating episodes in male mice. (a) Raster plot showing the temporal sequences in full and partial mating episode in male mice. Full mating (top, ejaculation achieved) and partial mating (bottom) episodes for 60 minutes following the introduction of a hormone-primed female mouse. Each row represents a single male mouse (full mating, n=13 mice; partial mating, n=11). Asterisks indicate sexually experienced males. (b) Representative photograph of male mouse (right) showing sniffing a female mouse (left) as appetitive behavior. (c) Representative photograph of male mouse (left) showing mount/intromission as consummatory behavior. (d, e) Temporal distribution of behavioral events (Sniffing, Mount, Intromission, Ejaculation, Digging) in the full (d) and partial (e) mating episodes. Each dot represents a single event. Vertical black bars indicate the median occurrence time. Sniffing occurring after ejaculation is not shown. (f) Time course of total interaction frequency of the full (blue, n=13 mice) and partial mating (red, n=11) groups. Shaded areas indicate s.e.m. (g) Sniffing frequency during the initial 15 minutes compared among the full mating (n=13), partial mating (n=11) episodes, and novel male introduction (n=12). Two-sided Welch’s *t-*test followed by Shaffer’s correction. (h) Number of USVs emitted during the first 1 minute of interaction in the full mating (n=11) and partial mating (n=7) episodes. Two-sided Welch’s *t-*test, N. S.: *P* = 0.207. Data are presented as mean ± s.e.m. \*\**P* < 0.01, \*\*\**P* < 0.001.

PM males initially engaged in active sniffing, but this declined over time (Figure 1e,1f, S1). The overall frequency of female-directed interactions in the PM group is comparable to that of the FM group (Figure 1f). PM males also showed a stronger investigatory response to females than to a male intruder, as indicated by a higher sniffing frequency during the first 15 minutes after introduction in a trial without overt aggression (Figure 1g). Consistent with previous reports that male courtship behavior is accompanied by ultrasonic vocalizations (USVs) ^15,16^, frequent USVs were detected during the first minute after female introduction in both FM and PM males (Figure 1h). Although some PM males attempted to mount, mounting and intromissions were infrequent and not sustained (Figure 1e, S1). Both sexually naïve and sexually experienced males were present in the PM, and sexually experienced males did not overrepresent in FM (p = 0.0995, Fisher’s exact probability test). These findings suggest that PM males retained appetitive sexual motivation, but showed a diminished propensity to engage in consummatory behaviors.

Digging behavior, displacing bedding material at multiple cage locations, was frequently observed after ejaculation in FM males, and was also present in PM males, most prominently in the latter half of the observation period (Figure S1b, S1e, S1h). Since digging is rare during a comparable observation window in singly housed males ^17,18^, it may reflect an internal state inversely related to sexual motivation, consistent with previous observations ^13^. Taken together, these findings suggest that C57BL/6J males may differ in internal state or drive, which determines whether they progress to consummatory behavior such as intromission or instead remain in the appetitive phase despite robust sniffing and USVs (Figure S1i, S1j).

### Consummatory mating preferentially recruits cMPOA *Esr1/Penk* neurons

Given that *Esr1*-positive cells in the MPOA can be composed of distinct subpopulations with different gene expression ^19,20^, we examined whether the activation of a specific subpopulation of *Esr1*-positive neurons is associated with the transition to consummatory behavior. After a 60-min mating behavior observation, the female was removed, 60 min later, brains were collected to assess c-Fos, an activation marker, with two rounds of isHCR for *Esr1*, *Penk*, *neurotensin*, *Vglut2*, and sexually dimorphic nucleus marker, *Moxd1* ^21^ (Figure 2a–2d). *Esr1*+/c-Fos+ cells were rarely observed under the home cage control condition. In both the *Esr1*-positive and -negative cell populations, many c-Fos-positive cells were observed in the FM mice that reached ejaculation, while the PM mice, which exhibited less consummatory behavior, showed intermediate values between the control and FM groups (Figure 2e). In FM males, both c-Fos+ and *Esr1*+ cells were distributed throughout the MPOA, with particularly high densities in the central MPOA (cMPOA) and the medial preoptic nucleus (MPN), consistent with a previous report ^22^. *Penk*+/*Esr1*+ cells were enriched in the cMPOA, whereas *neurotensin*+ cells were more prominent in the MPN. Most markers labeled non-overlapping populations (Figure 2d). Among *Esr1*+ cells, *Penk*+ cells in the cMPOA exhibited the highest number of c-Fos+ cells in FM males, showing a significant increase compared to both the home cage control and PM groups (Figure 2f). Among *Penk*/*Esr1* double-positive cells, 45.0% ± 7.2% were c-Fos-expressing neurons in the FM condition (Figure S2a). *Penk*+ neurons in the cMPOA had larger somata and were frequently *Esr1*+, whereas *Penk*+ neurons outside the cMPOA were smaller and mostly *Esr1-*, suggesting distinct neuronal populations (Figure S2e-S2g). *Neurotensin*+/c-Fos+ neuron numbers also increased in FM, compared to the home cage control and PM. However, the proportion of c-Fos+ cells among *neurotensin*/*Esr1* double-positive neurons was lower than that of *Penk+* neurons, with 18.1% ± 1.9% of *neurotensin*/*Esr1*+ neurons showing c-Fos expression in FM (Figure S2b). Among *Esr1*-negative cells, *Vglut2*+ cells showed more c-Fos+ cells in the FM than in home cage control and PM groups. Nevertheless, c-Fos-positive cells still accounted for fewer than 10% of *Esr1*–/*Vglut2*+ cells (Figure S2c). In *Moxd1*+ cells, the number of c-Fos+ cells did not differ between PM and FM, while the proportion of c-Fos + cells among *Moxd1*+ cells in FM was greater than that of PM (Figure 2f, S2d). A marker-negative population, putatively GABAergic, showed significantly higher c-Fos expression in both the PM and FM conditions than in controls, with no significant difference between PM and FM. Collectively, these findings identify cMPOA *Penk+* neurons as the largest *Esr1*-positive population showing c-Fos expression during consummatory mating.

**Figure 2.**
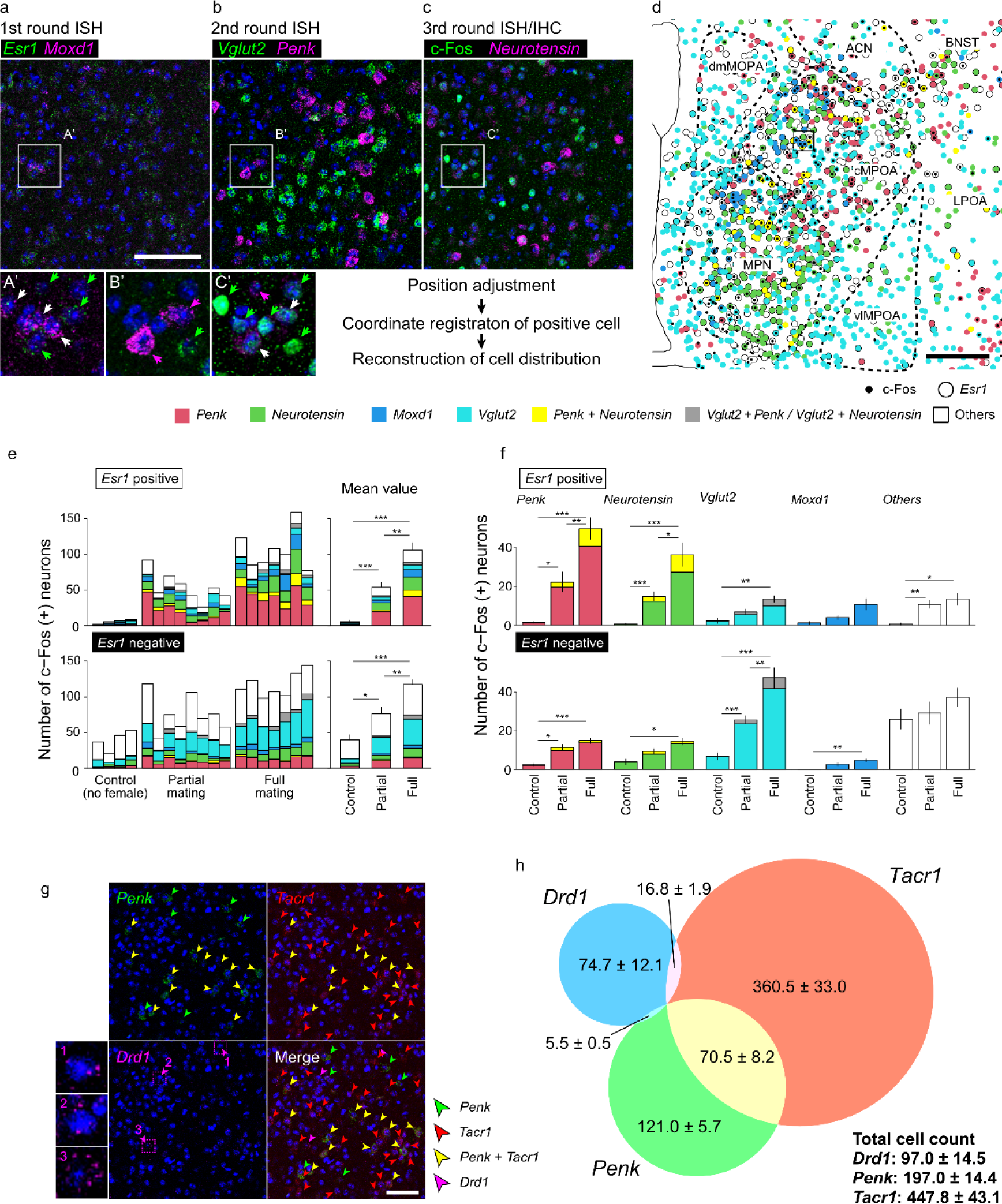
cMPOA Penk neurons are preferentially activated during consummatory sexual behavior. (a–c) Representative confocal images from three rounds of ISH-IHC, showing the same field of view. (a) First round ISH for *Esr1* (green) and *Moxd1*(magenta). (b) Second round ISH for *Vglut2* (green) and *Penk* (magenta). (c) Third round detection of c-Fos (green) and *neurotensin* (magenta). (a’-c’) Higher magnification views of the boxed regions in a-c. Nuclei were counterstained with Hoechst 33342. (d) Representative reconstruction showing the distribution of each cell subtype based on multi-round ISH-IHC data. Black dots indicate c-Fos-positive cells, and open circles denote *Esr1*-positive cells. Color coding: red, *Penk*; green, *neurotensin*; blue, *Moxd1*; light blue: *Vglut2*; yellow, *Penk* and *neurotensin*; gray, *Vglut2* and *Penk,* or *Vglut2* and *neurotensin*; white, others. ACN, anterior commissural nucleus; BNST, bed nucleus of the stria terminalis; cMPOA, central part of the medial preoptic area; dmMPOA, dorsomedial part of the medial preoptic area; LPOA, lateral preoptic area; MPN, medial preoptic nucleus; vlMPOA, ventrolateral part of the medial preoptic area. (e-f) Stacked bar graphs quantifying the numbers of c-Fos–positive cells classified by their gene expression profile under three behavioral conditions. Data are separated into *Esr1*-positive (top) and *Esr1*-negative (bottom) populations. Colors correspond to the gene markers defined in (d). The mean values within each group are shown in the right side (no female control: n = 4, partial mating: n = 8, full mating: n = 7). (f) Quantification of c-Fos expression in specific neuronal populations across three behavioral conditions. Upper panels show *Esr1*-positive population, and lower panels show *Esr1*-negative populations. Data are expressed as mean ± s.e.m. Two-sided Welch’s *t-*test followed by Shaffer’s correction. * *P* < 0.05, ** *P* < 0.01, *** *P* < 0.001. (g,h) Representative triple ISH images for *Penk*, *Tacr1*, and *Drd1* (g), and Venn diagram illustrating the cellular composition within the MPOA (h). (g1-g3) Magnified view of the boxed region in g. (h) Numbers indicate cell counts (mean ± s.e.m., n = 4 mice). Green, red, yellow, and magenta arrowheads indicate *Penk, Tacr1*, *Penk/Tacr1*, and *Drd1* positive cells, respectively. Nuclei were counterstained with Hoechst 33342. Scale bars: 100 µm (a-c), 200 µm (d), 50 µm (g).

Next, we examined whether MPOA *Penk+* neurons are distinct from neurons expressing *Tacr1* or *Drd1*, implicated in male sexual behavior ^13,14^. Triple-labeling for *Penk*, *Tacr1*, and *Drd1* mRNA revealed that about 35% of *Penk*+ cells were also *Tacr1*+, while 15% of *Tacr1*+ cells co-expressed *Penk* mRNA (Figure 2g, 2h). In contrast, *Drd1*+ cells were sparse in the MPOA, and *Drd1*/*Penk* double-positive cells were very few. Thus, in the MPOA, *Penk*+ neurons are largely distinct from *Drd1*+ neurons and also largely separate from *Tacr1*+ neurons.

### cMPOA Penk neurons show sustained Ca²⁺ dynamics during FM and encode key transitions

To investigate intracellular calcium dynamics of *Penk*+ neurons during male mating behavior, we performed fiber photometry in Penk-Cre males expressing calcium indicator jGCaMP8m in the cMPOA (Figure 3a, 3b). FM males displayed frequent calcium transients throughout the session, whereas PM males showed a brief response at initial female encounter followed by minimal activity (Figure 3c, 3d).

**Figure 3.**
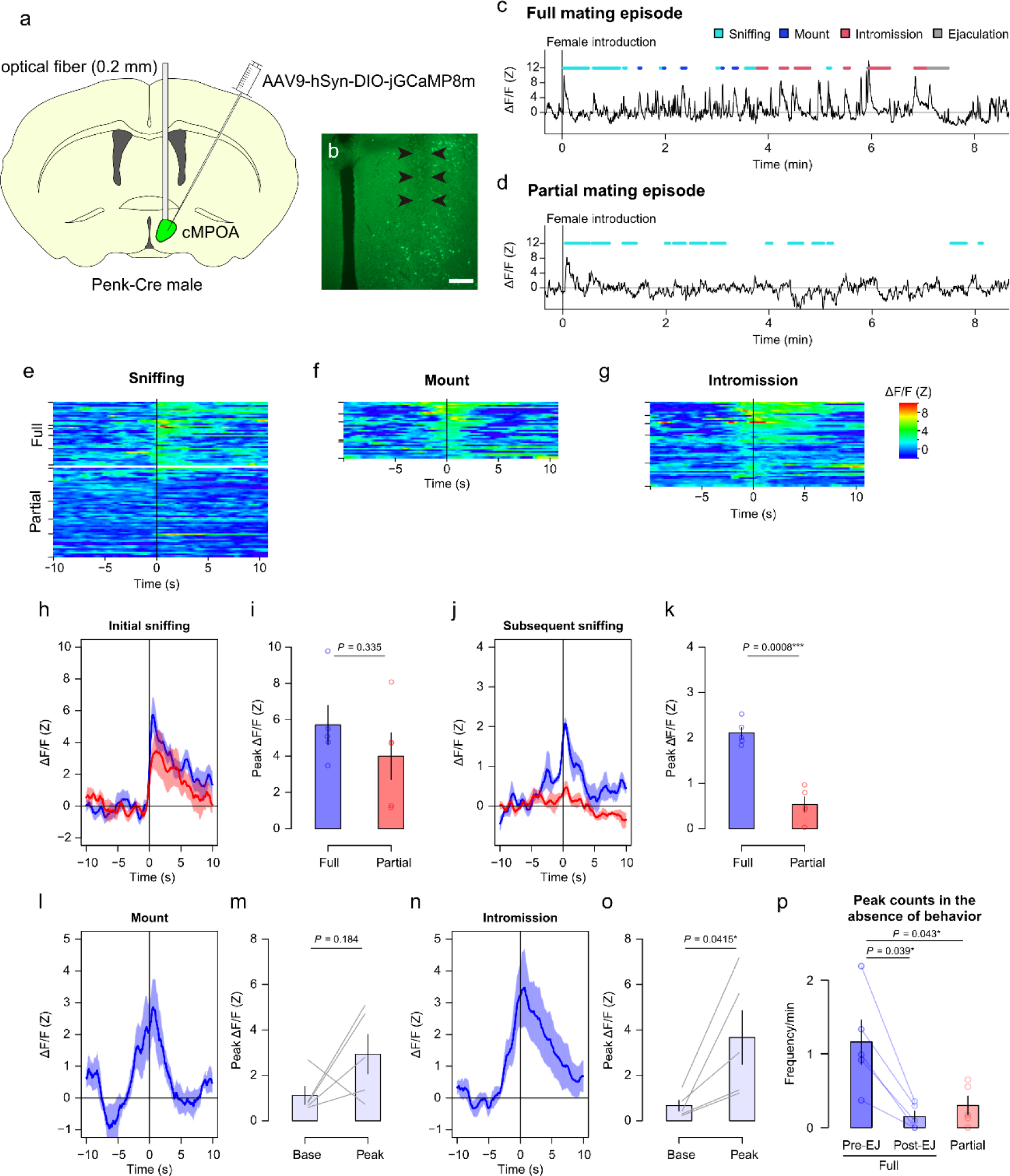
cMPOA Penk neurons exhibit time-locked activation during sexual consummatory phase. (a) Schematic diagram illustrating AAV injection to the cMPOA and implantation of an optical fiber positioned above the injection site in Penk-Cre male mice. (b) Representative fluorescence photomicrograph showing the optical fiber track (arrowheads) within the MPOA. Scale bar: 200 µm. (c, d) Representative ΔF/F traces during the full (c) and partial (d) mating episodes. Behavioral events are indicated by color-coded bars above the traces. Time is aligned to female introduction. (e-g) Heatmap of ΔF/F activity aligned to the onset of sniffing (e), mount (f), and intromission (g). In (e), traces are separated into the full (top) and partial (bottom) mating episodes. Each row represents a single behavioral event from one mouse (full mating: n = 5, and partial mating: n = 5). (h, i) ΔF/F traces (h) aligned to the onset of the first sniffing bout and peak ΔF/F values (i) for the full (blue) and partial (red) mating groups (5 mice per group). ΔF/F values were normalized by subtracting the median ΔF/F during -10s to -5s relative to onset. Two-sided Welch’s *t-*test. Shaded areas indicate s.e.m. for each frame. (j, k) ΔF/F traces (j) aligned to sniffing onset excluding the first sniffing bout, and peak ΔF/F values (k) for the full (blue) and partial (red) mating groups (5 mice per group). Two-sided Welch’s *t-*test. (l, m) ΔF/F traces (l) aligned to mount onset in the full mating group (5 mice per group). Baseline versus peak ΔF/F values associated with mounting were compared (m). Two-sided paired t-test. (n, o) ΔF/F traces (n) aligned to intromission onset in the full mating group (5 mice per group). Baseline versus peak ΔF/F values associated with intromission were compared (o). Two-sided paired t-test. (p) Frequency of peak events occurring outside annotated mating behaviors. Comparisons include the pre-ejaculation phase (Pre-EJ) and post-ejaculation phase (Post-EJ) within the full mating episodes, and the partial mating episodes. (full mating: n = 5, and partial mating: n = 5). Two-sided Welch’s *t-*test followed by Shaffer’s correction. Data are expressed as mean ± s.e.m.

In FM males, cMPOA-Penk+ neurons responded rapidly and consistently to sniffing. In PM males, the first sniffing bout elicited a calcium peak comparable to FM (Figure 3e, 3h, 3i), but subsequent sniffing bouts elicited little or no activity, with significantly reduced peak amplitudes relative to FM (Figure 3e, 3j, 3k). In FM, calcium signal rose and peaked at the onset of mounting and intromission (Figure 3f, 3g). However, peak intensity analysis showed that the mounting-associated increase in calcium signals did not reach statistical significance (Figure 3l, 3m). In contrast, a significantly elevated calcium peak was observed in the intromission (Figure 3n, 3o). Notably, FM males also exhibited frequent calcium peaks in the absence of overt behavior (Figure 3p, S3a–S3d). These behavior-independent peaks were significantly reduced after ejaculation, reaching levels comparable to PM males (Figure 3p). Control mice expressing Cre-dependent GFP did not show corresponding fluorescence changes (Figure S3e–S3j).

### Bidirectional chemogenetic control of cMPOA Penk neurons gates consummatory, not appetitive, behavior

To examine whether manipulation of cMPOA Penk+ neurons affects male sexual behavior, we chemogenetically activated cMPOA-Penk+ neurons by expressing Cre-dependent hM3Dq in Penk-Cre males (Figure 4a, 4b). Following intraperitoneal (i.p.) injection of either PBS or clozapine-n-oxide (CNO), a hormonally primed estrous female was introduced and male behaviors were recorded with USV measurements taken during the first minute (Figure 4c).

**Figure 4.**
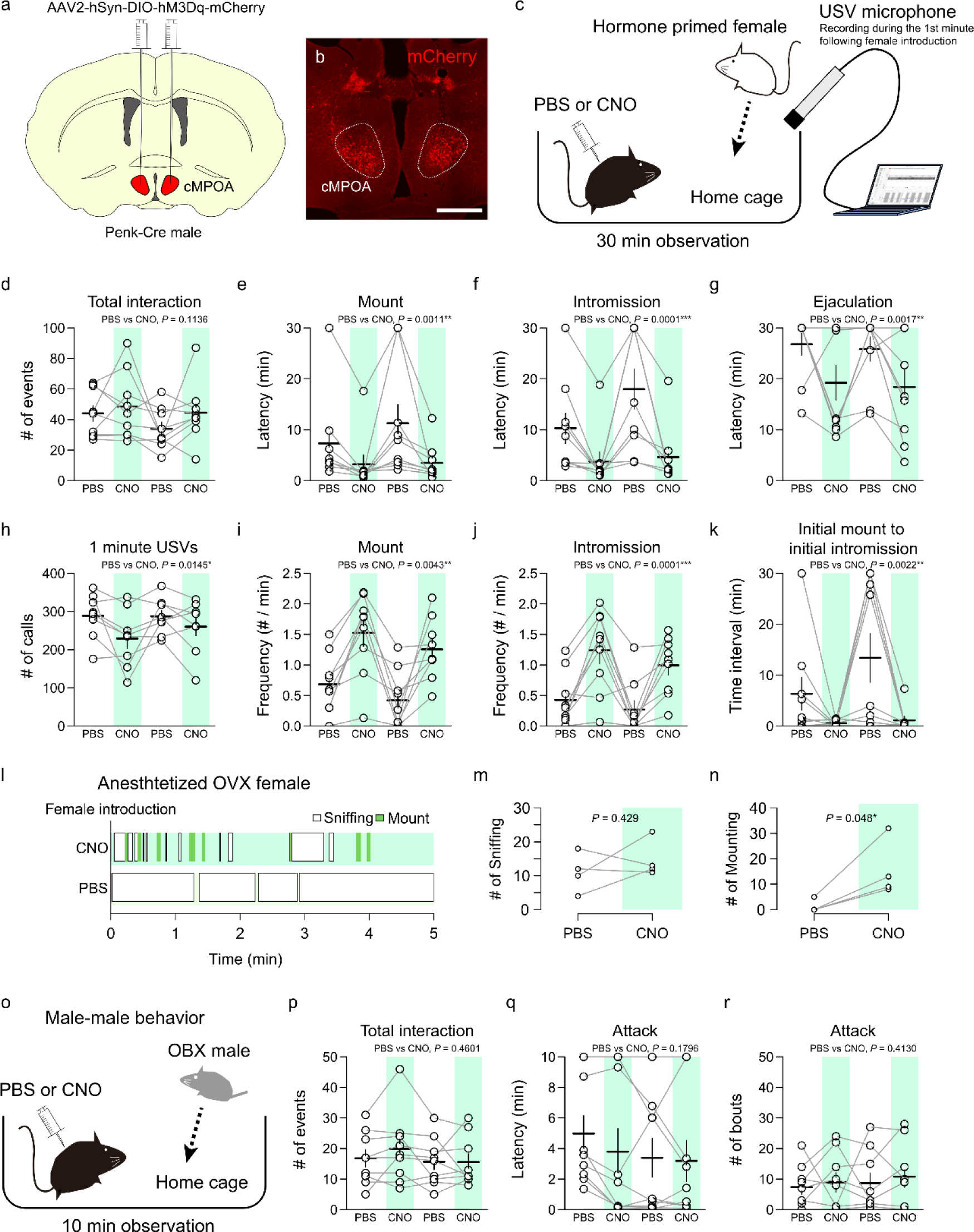
Chemogenetic activation of cMPOA Penk neurons facilitates male sexual behavior. (a) Schematic diagram illustrating AAV injection into the cMPOA of Penk-Cre male mice. (b) Representative fluorescence photomicrograph showing mCherry expression in the cMPOA. Scale bar: 500 µm. (c) Experimental scheme for the male-female mating assay. A hormone-primed female mouse was introduced to the home cage of a male mouse treated with either PBS or CNO (0.5 mg/kg) prior to the female introduction, and their behavior was observed for 30 minutes. USVs were recorded during the first minute after female introduction. (d–k) Quantification of male sexual behavior during a 30-minute observation period (9 mice). Lines connect observations from the same mice under PBS and CNO. Total number of interactions (d). Latency to the first occurrence of mount (e), intromission (f), and ejaculation (g). Number of USVs emitted during the first minute of interaction (h). Frequency of mounts (i) and intromissions (j). Interval from the first mount to the first intromission (k). Horizontal and vertical bars indicate means and s.e.m., respectively. Group differences were tested by type II ANOVA. (l–n) Mating behavior toward anesthetized ovariectomized (OVX) female mice (4 mice). (l) Representative raster plot showing the time course of sniffing (white boxes) and mounting (green boxes). (m) Number of sniffing events. Two-sided paired *t*-test, *P* = 0.429. (n) Number of mounts. Two-sided paired *t*-test, *P* = 0.048. (o) Experimental paradigm for the male–male resident–intruder test. An olfactory bulbectomized (OBX) male mouse was introduced to the home cage of a male mouse treated with either PBS or CNO prior to the intruder introduction, and their behavior was observed for 10 minutes. (p–r) Quantification of male–male interactions (9 mice). Total number of interactions (p), latency to the first attack (q), and number of attacks (r). Horizontal and vertical bars indicate means and s.e.m., respectively. Group differences were tested by type II ANOVA.

CNO administration markedly enhanced mounting and intromission, without significantly promoting appetitive behaviors, indicated by female-directed interactions and USVs (Figure 4d, 4h). CNO shortened latencies to mount and intromission and increased their frequencies (Figure 4e, 4f, 4i, 4j). The most prominent effect was a shortening of the interval from first mount to first intromission. In most trials, the first mount directly led to intromission or was followed within a few minutes (Figure 4k). CNO also shortened the latency to ejaculation (Figure 4g). After ejaculation, males exhibited a refractory period comparable to PBS controls, and no post-ejaculatory mounting was observed during 30-min observation. These effects were not observed in GFP-expressing controls (Figure S4a–S4i). In contrast, chemogenetic activation of cMPOA neurons in neurotensin-Cre mice did not affect any of the behavioral parameters examined (Figure S5a–S5h). In Vglut2-Cre mice, this manipulation reduced female-directed interactions and USVs but did not enhance consummatory behaviors (Figure S5i–S5p). These findings support a specific role for cMPOA-Penk+ neuron activation in facilitating consummatory behavior.

To minimize potential female-driven feedback, we introduced anesthetized ovariectomized (OVX) females (Figure 4l–4n). Although sniffing did not differ between PBS- and CNO-treated males, CNO administration enhanced mounting, suggesting that the activation of cMPOA-Penk+ neurons can directly drive male sexual behavior, independent of active female feedback.

We then examined the effect of cMPOA-Penk+ neuron activation on intermale behaviors. After the administration of PBS or CNO, an olfactory bulbectomized (OBX) male was introduced into the home cage of the test mouse (Figure 4o). Because OBX mice are less aggressive, using an OBX mouse as the intruder reduces intruder-initiated interactions toward the test mouse and thereby decreases variability in the test mouse’s aggressive behavior. CNO treatment did not affect male-directed interactions, attack latency, or attack frequency (Figure 4o–4r), and mounting was not observed. CNO administration to mice expressing GFP showed no changes in aggressive behavior (Figure S4j–S4l), except for attack latency, likely reflecting experience effects from repeated tests (Figure S4k). Thus, cMPOA-Penk+ neuron activation promotes sexual behavior in a female-directed context and does not induce male-directed behaviors, which was not observed when *Esr1*+ or *Tacr1*+ neurons in the MPOA were activated ^11,14^.

Next, we suppressed cMPOA-Penk+ neurons using Cre-dependent hM4Di (Figure 5a, 5b). Thirty minutes after PBS or CNO administration, females were introduced. CNO administration reduced the total number of interactions toward the female (Figure 5c, 5d), increased sniffing counts (Figure 5e), and did not affect USVs immediately after female introduction (Figure 5h). In contrast to appetitive behaviors, mounting and intromission were strongly suppressed by CNO (Figure 5g–5j) and reduced ejaculation within 30 minutes (1/9 CNO vs. 8/9 PBS, *p* = 0.0034, Fisher’s exact probability test) (Figure 5l). Vaginal plugs were nonetheless detected in all females the next day, consistent with delayed ejaculation as CNO effects waned. Notably, CNO treatment also enhanced digging (Figure 5m, 5n). These findings suggest that suppression of cMPOA-Penk+ neurons preferentially impairs consummatory behaviors while sparing appetitive responses, producing a PM-like phenotype.

**Figure 5.**
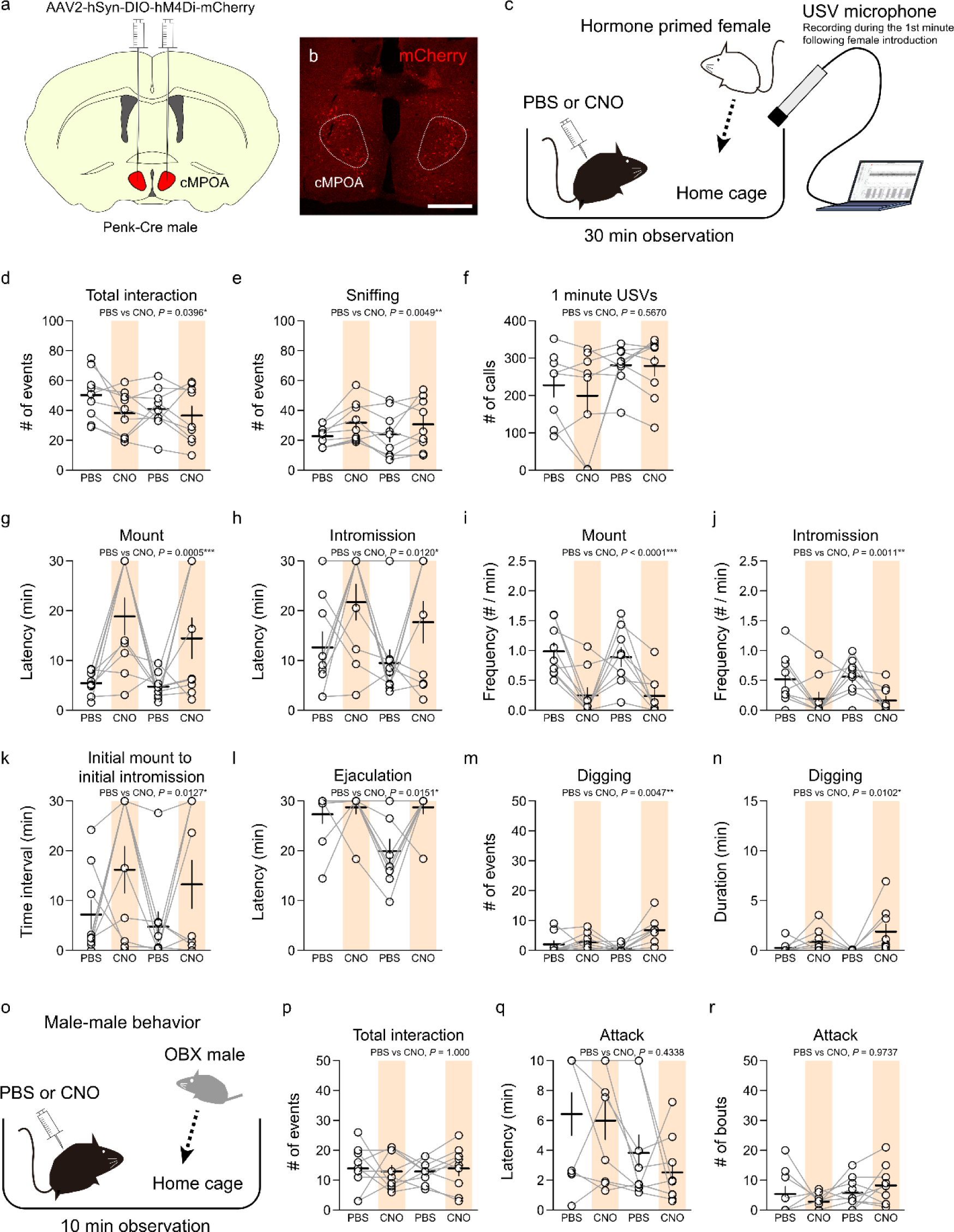
Silencing cMPOA Penk neurons suppresses male sexual behavior. (a) Schematic diagram illustrating AAV injection into the cMPOA of Penk-Cre male mice. (b) Representative fluorescence photomicrograph showing mCherry expression in the cMPOA. Scale bar: 500 µm. (c) Experimental scheme for male-female mating assay. A hormone-primed female mouse was introduced to the home cage of a male mouse treated with either PBS or CNO (1.25 mg/kg) prior to the female introduction, and their behavior was observed for 30 minutes. USVs were recorded by USV microphone during the first minute following female introduction. (d–n) Quantification of male sexual behaviors during a 30-minute observation period (9 mice). Lines connect observations from the same mice under PBS and CNO. Total number of interactions (d), sniffing (e), and number of USV calls during the first minute of interaction (f). Latency to the first occurrence of mount (g) and intromission (h). Frequency of mounts (i) and intromissions (j). Interval from the first mount to the first intromission (k). Latency to ejaculation (l). Number (m) and duration (n) of digging behaviors. Horizontal and vertical bars indicate means and s.e.m., respectively. Group differences were tested by type II ANOVA. (o) Experimental paradigm for the male–male resident–intruder test. An olfactory bulbectomized (OBX) male mouse was introduced to the home cage of a male mouse treated with either PBS or CNO prior to the intruder introduction, and their behavior was observed for 10 minutes. (p–r) Quantification of male–male interactions (9 mice). Total number of interactions (p), latency to the first attack (q), and number of attacks (r). Horizontal and vertical bars indicate means and s.e.m., respectively. Group differences were tested by type II ANOVA.

As with activation, inhibition of cMPOA-Penk+ neurons did not alter intermale behaviors in intruder assay using OBX males (Figure 5o). CNO administration had little effect on male-directed interactions, attack latency, attack number, and neither mounting nor digging was observed (Figure 5p–5r).

### Optogenetic stimulation of cMPOA-Penk+ neurons reveals a slow-time scale facilitation of consummatory behaviors

To refine temporal requirements, we expressed Cre-dependent ChR2 in the cMPOA of Penk-Cre mice (Figure 6a, 6b). In response to light stimulation under basal conditions, approximately 70% of ChR2-expressing neurons expressed c-Fos, which was higher than mice expressing GFP (Figure 6c, 6d).

**Figure 6.**
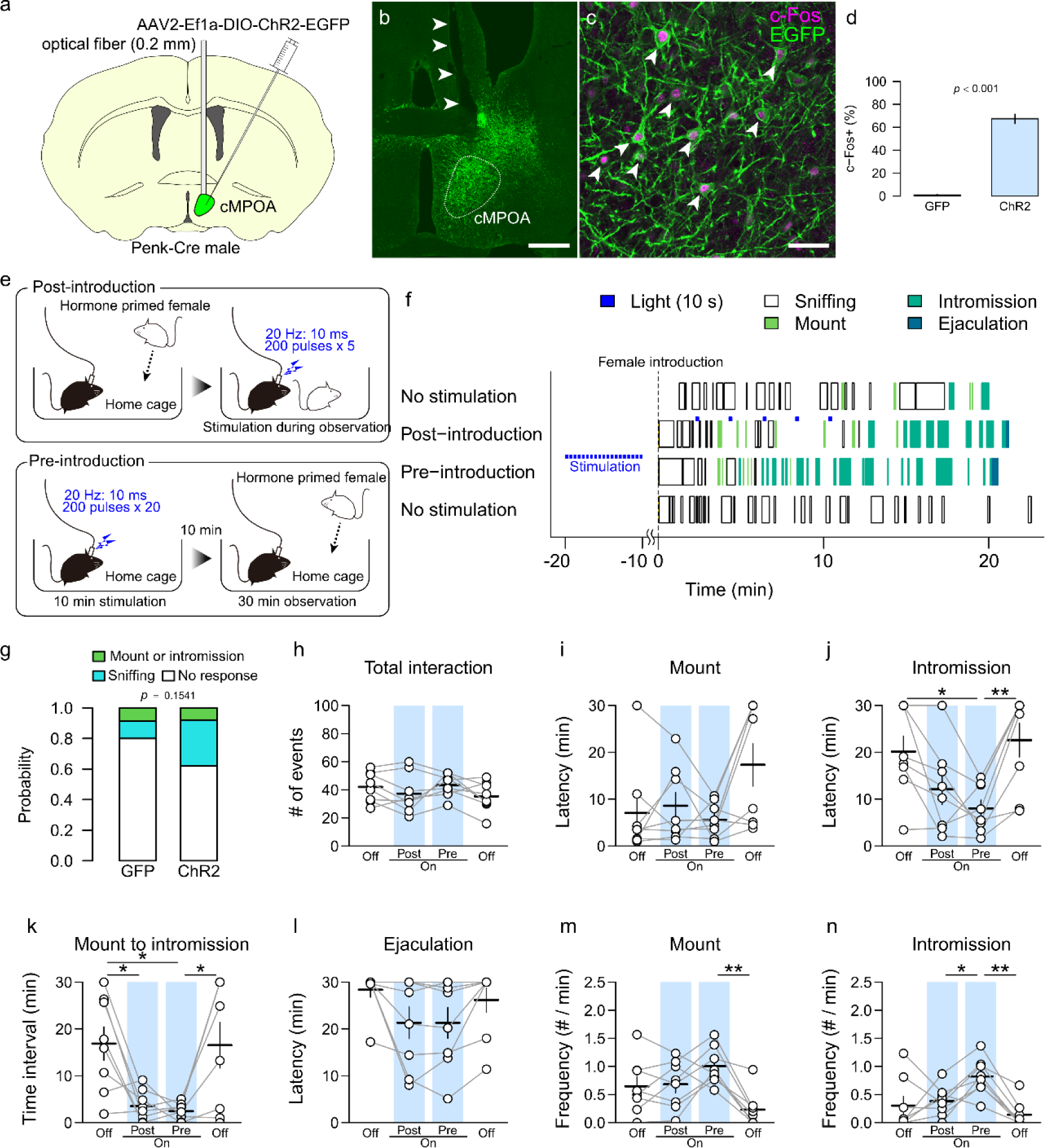
Optogenetic activation of cMPOA Penk neurons induces a persistent state of sexual arousal. (a) Schematic drawings illustrating the injection of AAV2-Ef1α-DIO-ChR2-EGFP or AAV-Ef1α-DIO-EGFP into the cMPOA and the implantation of an optical fiber positioned above the injection site in Penk-Cre male mice. (b) Representative fluorescence photomicrograph showing the optical fiber track (arrowheads) within the cMPOA. (c, d) Validation of optogenetics activation observed in ChR2-expressing cells after light stimulation. Representative immunofluorescence image (c) c-Fos staining following photostimulation. Arrowheads indicate cells positive for c-Fos and GFP. Percentage of c-Fos-positive cells in GFP-positive cells (d). 7 mice per group. Two-sided Welch’s *t*-test. *P* < 0.001. (e) Experimental paradigm for the Pre/Post-introduction stimulation. In the post-introduction condition, a female mouse was introduced into the male’s home cage, and optical stimulation (20 Hz, 10 ms pulses, 200 pulses per train) was delivered once every 2 minutes, repeated 5 times. In the pre-introduction condition, beginning 20 minutes before the introduction of the female, the same optical stimulation (20 Hz, 10 ms pulses, 200 pulses) was delivered once every 30 seconds, repeated 20 times. (f) Representative raster plot showing the time course of male sexual behavior aligned to female introduction. Photostimulations are indicated by blue marks. (g) Probability of immediate behavioral responses to photostimulation in the post-introduction condition. No significant difference between GFP (35 events from 7 mice) and ChR2 mice (37 events from 8 mice). Fisher’s exact probability test. *P* = 0.1541. (h–n) Quantitation of male sexual behavior under different stimulation conditions (n = 8 mice). Lines connect observations from the same mice. Total number of interactions (h). Latency to the first occurrence of mount (i) and intromission (j). Interval from the first mount to the first intromission (k). Latency to ejaculation (l). Frequency of mounts (m) and intromissions (n). Significant differences are indicated by asterisks. Paired *t*-test followed by Shaffer’s correction. * *P* < 0.05, ** *P* < 0.01. Horizontal and vertical bars indicate means and s.e.m., respectively. Data are expressed as mean ± s.e.m. Scale bars: 500 µm (b) and 50 µm (c).

We first tested whether brief stimulation could acutely trigger sexual behaviors. When light was delivered five times at 2-minute intervals after female introduction (“post-introduction” stimulation, Figure 6e), male sexual behavior did not change immediately after stimulation, and 60% of trials showed no female-directed behavior (Figure 6f, 6g). Mounting was occasionally observed but its frequency was comparable to controls (Figure 6g). Nonetheless, the interval from first mount to first intromission was significantly shortened (Figure 6k), although intromission rarely occurred during or immediately after stimulation. These observations suggest that activation of cMPOA-Penk+ neurons does not acutely trigger sexual behavior, but can facilitate the transition from mounting to intromission.

Next we tested whether transient activation of cMPOA-Penk+ neurons prior to female exposure could produce a longer-lasting facilitation of male sexual behavior. Delivering light pulses every 30 seconds for 20 total pulses, starting 20 minutes before female introduction and ending 10 minutes before introduction (“pre-introduction” stimulation, Figure 6e) enhanced sexual behavior after female introduction (Figure 6f). While total interaction counts and mount latency were unchanged, intromission latency, the mount-to-intromission interval, and the frequencies of mounting and intromission were all significantly altered compared to the no-stimulation control (Figure 6i–6n). These effects were not observed in GFP-expressing controls (Figure S6). Thus, cMPOA-Penk+ neuron activation increases the probability of entering consummatory sequences over a timescale of tens of minutes rather than acting as an immediate trigger.

### cMPOA Penk+ projections to VTA and PAG differentially promote mounting and intromission

To map downstream of cMPOA-Penk+ neurons, we visualized projections of cMPOA-Penk+ neurons using Cre-dependent, palmitoylated EGFP in Penk-Cre mice (Figure S7a, S7b). Dense GFP-positive fibers were observed in the paraventricular nucleus (PVN), dorsomedial hypothalamus (DMH), ventromedial hypothalamus (VMH), tuberomammillary nucleus (TMN), posterior hypothalamus (PH), ventral tegmental area (VTA), periaqueductal gray (PAG), and Barrington’s nucleus (BAR) (Figure S7c–S7h), whereas fibers were sparse in the principal nucleus of the bed nucleus of the stria terminalis and the medial amygdala, regions that have previously been implicated for sexual behavior ^23–25^.

Given the established roles for VTA and PAG in reward and motor execution, respectively ^5,14,26^, we optogenetically stimulated cMPOA-Penk+ axon terminals in each region using a pre-introduction paradigm (Figure 7a,7h). VTA terminal stimulation reduced mount latency and increased mounting frequency (Figure 7d, 7f) without significant changes in intromission latency or frequency, nor the mount-to-intromission interval (Figure 7c, 7e, 7g), consistent with a selective effect of VTA on mounting, but not on intromission ^14^. In contrast, PAG terminal stimulation enhanced both mounting and intromission. PAG stimulation reduced the latencies to both behaviors, shortened the mount-to-intromission interval, and typically yielded intromission within 5 minutes of the first mount (Figure 7j–7l). PAG terminal stimulation also increased mounting and intromission frequencies and total social interactions (Figure 7i, 7m, 7n). Together, these findings suggest that the projections from cMPOA-Penk+ neurons to the VTA enhance motivation for mounting, whereas the projections to the PAG facilitate execution and the transition to intromission, and demonstrate that sustained facilitation by cMPOA-Penk+stimulation can be recapitulated by terminal stimulation.

**Figure 7.**
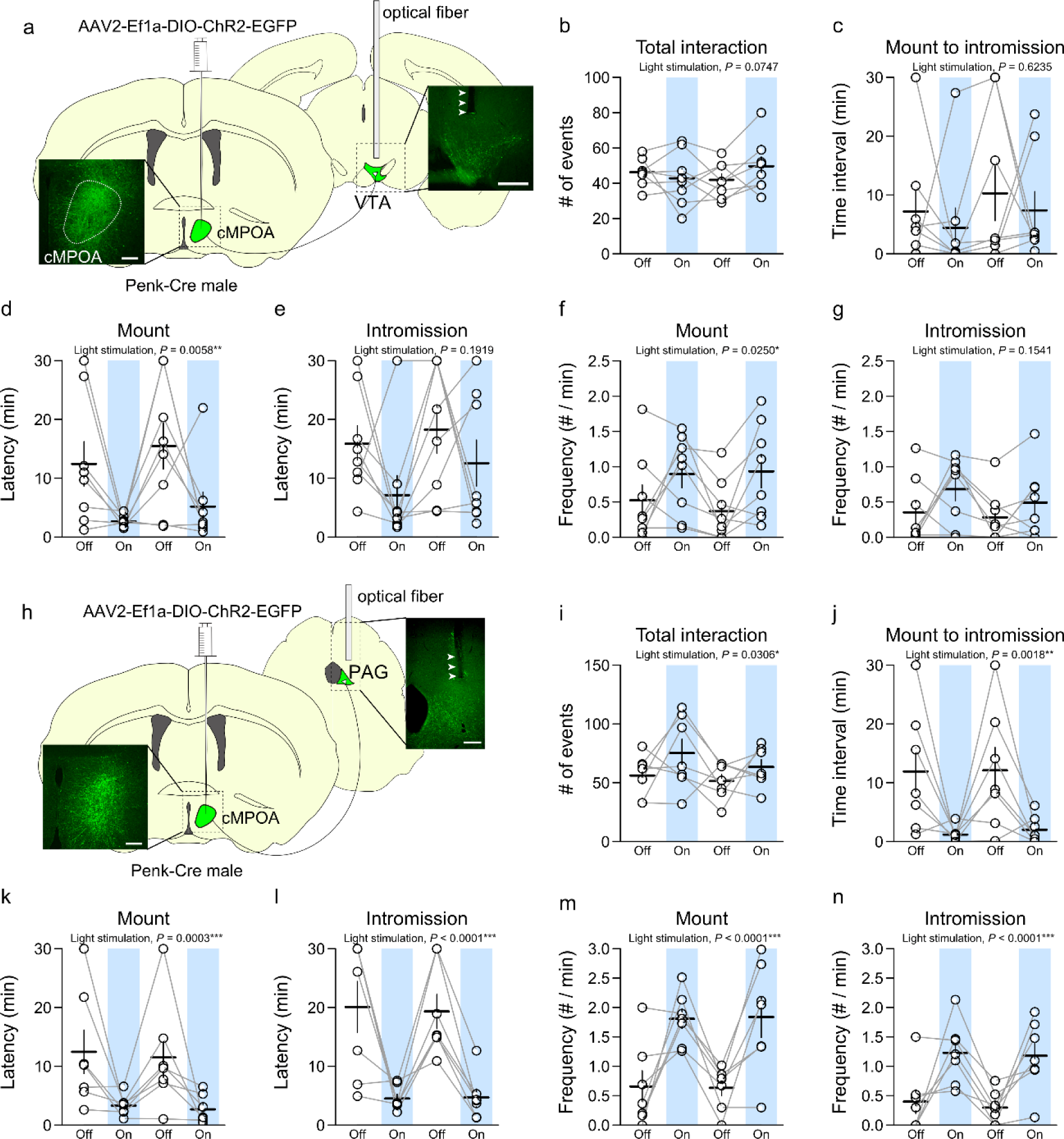
cMPOA Penk neurons project to the PAG to regulate consummatory sexual behavior. (a) Schematic illustration of viral tracing and optogenetic terminal stimulation. AAV2-Ef1α-DIO-ChR2-EGFP was injected into the cMPOA and optical fibers were implanted bilaterally above the VTA of Penk-Cre mice. Representative fluorescence images showing GFP-positive cells in the cMPOA and GFP-positive fibers in the VTA with optical fiber tracks (arrowheads). (b–g) Quantitation of male sexual behavior in response to optogenetic stimulation of the VTA (n = 8 mice). Total number of interactions (b). Interval from the first mount to the first intromission (c). Latency to the first occurrence of mount (d) and intromission (e). Frequency of mounts (f) and intromissions (g). (h) Schematic illustration of the viral tracing and optogenetic terminal stimulation. AAV2-Ef1α-DIO-ChR2-EGFP was injected into the cMPOA, and optical fibers were implanted bilaterally above the PAG in Penk-Cre male mice. Representative fluorescence images showing GFP-positive cells in the cMPOA and GFP-positive fibers in the PAG with optical fiber tacks (arrowheads). (i–n) Quantitation of male sexual behavior in response to optogenetic stimulation of the PAG (n = 7 mice). Total number of interactions (i). Interval from the first mount to the first intromission (j). Latency to the first occurrence of mount (k) and intromission (l). Frequency of mounts (m) and intromissions (n). Horizontal and vertical bars indicate means and s.e.m., respectively. Group differences were tested by type II ANOVA.

## Discussion

The present study shows that activation of cMPOA-Penk+ neurons is associated with the completion of consummatory behavior. These neurons exhibited enhanced calcium activity from the time of female contact until ejaculation. Moreover, brief activation of cMPOA-Penk+ neurons before female introduction promoted progression of mating behavior to ejaculation, suggesting that cMPOA-Penk+ neurons play a crucial role in facilitating progression toward copulatory completion.

Unlike MPOA *Tacr1* neurons, which can trigger mounting in a command-like manner toward males and toys ^14^, cMPOA-Penk neuron activation did not immediately elicit mounting. Instead, it produced a sustained facilitating effect lasting on the order of ∼ 10 minutes, promoting progression across the consummatory phase. Activation selectively enhanced female-directed behaviors and was followed by a post-ejaculatory refractory period, consistent with modulation of physiologically typical mating patterns in C57BL/6J mice. Together with the spontaneous calcium activity observed beginning upon exposure to a female, these results suggest that cMPOA-Penk+ neuron activity reflects an internal state signal, consistent with sexual drive/arousal, that lowers the threshold for behavioral expression and supports progression toward the completion of copulation.

Chemogenetic activation of cMPOA-Penk+ neurons did not increase early social interaction or USVs immediately after female introduction. Because USVs are regarded as a precopulatory behavior that reflects appetitive motivation in male mice ^16,27–29^, this pattern supports the interpretation that cMPOA-Penk+ neurons primarily modulate processes downstream of appetitive motivation. Similarly, behavioral changes following chemogenetic inhibition were largely restricted to mounting and intromission. Notably, inhibition increased digging behavior, resembling profiles in PM mice. In contrast, stimulation of *Esr1*+/*Vgat*+ or *Tacr1*+ neurons can elicit USV production and male-directed mounting, and can strongly suppress aggression toward other males ^11,12,14^.

The MPOA is well established as a hypothalamic hub for male sexual behavior ^30–32^, but it comprises diverse neuronal subtypes (Tsuneoka, Yoshida, et al., 2017; Moffitt et al., 2018). *Esr1*-positive neurons in the MPOA are critical for both male sexual behavior and parental behavior ^10,33–37^. It is suggested that the *Esr1*-positive populations are functionally heterogeneous and can be subdivided by marker genes including *Vglut2*, *Penk*, *neurotensin*, *Moxd1*, *Drd1*, and *Tacr1*. Indeed, most *Penk*+ neurons in the cMPOA are *Esr1*-positive. cMPOA-*Penk*+ neurons were largely distinct from *Drd1*+ MPOA neurons and only partially overlapped *Tacr1*+ cells. Thus, cMPOA-Penk+ neurons are a novel Esr1 subgroup that regulates male sexual behavior in a manner distinct from *Drd1*+ or *Tacr1*+ neurons implicated in partner recognition or the motor execution of copulatory acts ^13,14^. Notably, elevated cMPOA-Penk+ activity exerted sustained effects over a behaviorally relevant timescale and promoted sexual behavior.

We further demonstrate that the projections from cMPOA-Penk+ neurons to the PAG and VTA contribute to the promotion of consummatory behavior as shown by terminal stimulation delivered 10 minutes before female introduction. Projection-specific stimulation suggests separable downstream modules: VTA terminal stimulation preferentially enhanced mounting without reliably improving intromission metrics, whereas PAG terminal stimulation promoted both mounting and intromission and robustly shortened the mount-to-intromission interval. This division is consistent with broader evidence that midbrain reward/motivation systems and brainstem motor-pattern circuits differentially support sexual performance and copulatory reflexes, and supports a model in which cMPOA Penk+ neurons broadcast a state signal that can be read out by distinct effectors to promote behavioral completion.

The persistence of behavioral facilitation after brief pre-introduction stimulation, together with sustained and behavior-independent calcium events during FM, is consistent with *Penk*+ neurons engaging slow neuromodulatory mechanisms, such as neuropeptide release, that gradually reshape excitability and downstream circuit gain. Importantly, the finding that optogenetic stimulation of fiber terminals alone can produce a sustained facilitative effect is consistent with this model. Although this study does not directly test opioid receptor involvement, the Penk identity provides a mechanistic hypothesis: enkephalinergic signaling in the MPOA-to-PAG/VTA pathways may implement an accumulator-like process that promotes copulatory completion over tens of minutes. Further mechanistic analysis will be necessary to test this possibility.

The internal changes that facilitate consummatory behaviors under the control of cMPOA-Penk+ neurons may reflect an increased sexual arousal. Although sexual arousal is widely accepted, its precise nature remains unclear. Sexual arousal is generally considered distinct from sexual motivation and refers to specific physiological and psychological responses to sexual stimuli ^38–41^. It is often proposed that arousal accumulates to enable intromission and ejaculation, supporting the progression of copulatory behavior. Several studies link MPOA activity to sexual arousal ^32,42,43^, but underlying mechanisms remain unknown. The cumulative, sustained effects of cMPOA-Penk+ neuronal activity, its relative independence from appetitive motivation, and its facilitation of consummatory behavior, particularly the shortening of intromission latency, closely match features typically attributed to sexual arousal.

In summary, cMPOA-Penk+ neurons may broadcast a sustained internal state signal, probably linked to sexual arousal, via PAG and VTA, that lowers behavioral thresholds and promotes efficient progression through consummatory mating to ejaculation. These findings support a modular MPOA architecture in which distinct Esr1 subtypes separately mediate rapid action initiation and slower state-driven facilitation of completion.

## Materials and Methods

### Animals

Adult mice 10-24 weeks of age were used for all experiments. Original breeding pairs of C57BL/6J mice and DBA2/J were obtained from Japan SLC and CLEA Japan. Male mice of Penk-Cre, neurotensin-Cre and Vglut2-Cre strains were obtained from JAX. Cre mice backcrossed to the C57BL/6J strain for more than ten generations were used. Mice were bred and maintained in our colony under controlled conditions (12 h light/dark cycle, lights on at 8:00 A.M., 23 ± 2 °C, 55 ± 5% humidity, and ad libitum access to water and food). Mice were weaned at 4 weeks of age and maintained in group housing until a week before surgery. Penk-Cre mice were transferred to a different room with light-off scheduled at 2:00 PM one week before surgery and were housed individually thereafter. Unless otherwise specified, male mice were paired once per week with hormone-primed females overnight beginning one month after surgery to train sexual behavior. This training continued weekly until two copulatory plugs were confirmed, thereby standardizing sexual experience. Males in which plugs were not observed for three consecutive sessions were classified as non-copulating males ^44^ and excluded from subsequent experiments. Female C57BL/6J mice employed as sexual behavior partners and male DBA/2J mice used for intermale aggression tests were likewise transferred to the same room one week before surgery. Female mice were subjected to ovariectomy under isoflurane anesthesia. OVX-females were subcutaneously injected with 0.02 mg β-estradiol 2 days and 1 day before the behavioral experiment, and were administered 0.3 mg progesterone 6 h prior to the experiment. Male DBA/2J mice were anesthetized with isoflurane, and olfactory bulbs were aspirated. Beginning one month after surgery, these mice were temporarily introduced at least twice into the cages of stud males. In females, it was confirmed that hormone priming induced sexual behavior within several hours, whereas in DBA/2J males, aggressive behavior was confirmed within 10 minutes after introduction. After confirming that these behaviors were induced, the mice were subsequently used multiple times in random order, while experiments were conducted to avoid repeated pairings of the same mice. All animal procedures were conducted in accordance with the Guidelines for Animal Experiments of Toho University and were approved by the Institutional Animal Care and Use Committee of Toho University (Approval Protocol ID #18-55-224, 19-51-405, 24-557).

### Viruses

The recombinant plasmids used for AAV production, pAAV-hSyn-DIO-hM3Dq-mcherry, pAAV-hSyn-DIO-hM4Di-mcherry were kind gifts from Dr. Brian Roth, pAAV-Ef1a-DIO-ChR2-EGFP and pAAV-Ef1a-DIO-EGFP were obtained from Addgene (#35507, 37084), and pAAV-Ef1a-DIO-palEGFP were kind gift from Dr. Daiki Nakatsuka ^45^. Other AAV packaging plasmids, pHelper and pACG2-y730f, were obtained from TAKARA (TKR 6230) and National Gene Vector Biorepository, respectively. The tsA201 cells were cultured in Dulbecco’s modified Eagle medium (DMEM) containing high glucose, glutamine, and pyruvate, supplemented with 10% Fetal Clone II serum and penicillin–streptomycin, until they reached a state without contact inhibition. The cells were then transfected with plasmids for AAV packaging using linear polyethylene imine (PEI-MAX, Polyscience). Three days after transfection, cells were detached using a scraper, centrifuged, and the resulting pellet was collected. Viral purification was performed by the previously described method with some modifications ^46^. After three freeze–thaw cycles to destroy the cell membranes, the samples were treated with DNase I (25U/ml, 2270A Takara) and RNase A (0.1mg/ml, R6148, Sigma) at 37°C 30 min. Cell membranes were further destroyed by incubation with sodium deoxycholate at a final concentration of 0.5% at 37 °C for 60 min. The cell lysate was then centrifuged at 15,000 rpm for 5 min at room temperature, and the supernatant was collected. The supernatant was subjected to chloroform extraction, and the aqueous phase was recovered. The recovered aqueous phase was precipitated with 8% PEG8000 and 0.5 N NaCl, followed by centrifugation to collect the pellet. The pellet was resuspended in HEPES buffer (pH 7.2) and subjected to an additional chloroform purification. The aqueous phase was then subjected to phase extraction using 10% PEG8000 and 13.5% ammonium sulfate, and the lower phase was collected. The recovered viral solution was concentrated using a 50-kDa ultrafiltration filter (Amicon Ultra, UFC9050, Millipore). The concentrated virus was buffer-exchanged into artificial cerebrospinal fluid and stored at −80 °C until use. Viral titers were confirmed to be more than 10^12^ copies/ml by quantitative PCR. Purified AAV for hSyn-DIO-jGCaMP8m (Addgene viral prep #162378-AAV9; RRID:Addgene_162378) was obtained from Addgene (deposited by the GENIE Project) ^47^.

### Stereotaxic surgeries

Surgeries on mice were performed under isoflurane inhalation anesthesia. Viruses were delivered into male mice aged 11–14 weeks using a stereotaxic apparatus (SR-5M, Narishige) and glass capillaries under oil pressure. Injections were targeted to the cMPOA at the following coordinates: AP +0.1 mm, lateral 0.5 mm, ventral 5.2 mm from bregma. For optogenetics, tracing and fiber photometry studies, 150 nL of virus was injected unilaterally into the brain. In chemogenetic studies, 250 nL of virus was injected bilaterally. Viruses were infused at a rate of 50 nL/min using a syringe holder (Narishige) and hand-made syringe pump processed by Arduino, and the needle was left for an additional 5 min and withdrawn gently. In mice used for optogenetics and fiber photometry, optical fibers (200 μm diameter, 0.39 NA, FT200EMT, Thorlabs) were stereotaxically implanted following viral delivery. Optical fibers were secured to the skull with dental cement on a base constructed using a 3D printer. In optogenetics experiments, optical fibers were implanted 0.5mm dorsal to the target coordinates, whereas in fiber photometry experiments, fibers were implanted 0.2mm dorsal to the target. In experiments involving optogenetic stimulation of VTA or PAG terminals, optical fibers were implanted at the following coordinates: VTA, AP −3.1 mm, lateral 0.5 mm, ventral 3.8 mm; PAG, AP −4.4 mm, lateral 0.5 mm, ventral 1.5 mm. Off-target samples in which viral injection sites were misplaced and more than half of the infection were located outside the cMPOA were excluded from the analysis, or samples with evidently few infected cells, were excluded from the analysis.

### Fiber photometry

Implanted optical fibers were secured to a 3D-printed baseplate, which can be directly connected to a modified version of the custom miniature fluorescence microscopy system ^48^. The miniature fluorescence microscope was made using a 3D printer, and contained an objective lens (achromatic lens, 4 mm diameter, 6 mm focal length; 63-690, Edmund Optics) and an imaging lens (achromatic lens, 4 mm diameter, 10 mm focal length; 64-519, Edmund Optics) positioned at the bottom. Above these, an EGFP fluorescence filter set (49002, Chroma), cut to a width of 4 mm, was placed. A blue LED (465–485 nm, Xlamp XQ-E, CREE) driven at 90 mA under constant current control was located laterally, and a miniature CMOS sensor (OVM6946, COMedia Ltd.) was positioned on the side opposite to the lens assembly. Video signals were obtained at 30 fps processed by a backend module (C8296, COMedia Ltd.) and recorded as a movie.

To habituate mice to the miniature microscope, dummy microscopes and cables were attached beginning one month prior to experiments, and two sessions of sexual behavior training were conducted under this condition. About 6 hours before behavioral testing, the dummy was replaced with the miniature microscope, and cables were connected to the computer a few minutes before the start of the experiment. Video for fluorescent recording and behavioral video recordings were synchronized by pulse currents generated by an Arduino controlling the LED power supply. Video files imaging the fiber tip were processed in ImageJ to extract only the cross-sectional images of the optical fiber. The brightness of regions outside the fiber cross-section was used as background signal, and the difference was defined as the fluorescence intensity. Data were excluded from subsequent analyses if the tip of the optical fiber moved within the video image or if frequent image noise was detected.

Raw fluorescence data were denoised by applying a moving average with a 20-frame (0.66-s) sliding window. Baseline fluorescence was determined using locally estimated scatterplot smoothing (LOESS) applied to data excluding the period of sexual behavior events and the 5-s periods before and after each event. The baseline was then subtracted from the raw fluorescence data and divided, yielding the instantaneous ΔF/F values. Z score normalized ΔF/F was calculated as (ΔF/F – mean (ΔF/Fbase)) / std (ΔF/Fbase), where ΔF/Fbase refers to the ΔF/F signal during the 2-min period before female introduction. For peri-event time plot (PETP) analysis, time zero was defined as the onset of the behavioral or target event. The median fluorescence value within the 10- to 5-s window preceding the event was used as a normalization factor, and changes in fluorescence from baseline were calculated as acute response during a behavior. For amplitude change analysis around events, the 95% peak fluorescence value within the 5-s window of baseline and after each event was calculated and compared. The data were excluded from analysis if there was an overlap of other behaviors within this time window. The average response of all behavioral event for each episode was then calculated for population analysis.

For peak frequency analysis, a peak was defined as a local maximum with a Z value greater than 3 that persisted for at least 0.1 seconds, showing an increase of at least 3 above the median of the preceding 1 second, with a minimum inter-peak interval of 0.5 seconds.

### Histology

Mice were deeply anesthetized with isoflurane and perfused with 4% PFA in PBS. Brains were removed and post-fixed overnight at 4 °C in 4% paraformaldehyde (PFA)/PBS, followed by cryoprotection in 30% sucrose/PBS solution at 4 °C for 2 days. Brains were cryo-embedded in Surgipath FSC22 and stored at −80 °C until sectioning. For three-round multiplex ISH/IHC staining, brains were sectioned at 10 μm, and every 12th section was mounted on glass slides and stored at −20 °C until staining. For other staining procedures, brains were cryo-sectioned at 40 μm, and every 3rd section was collected in cryoprotectant solution (1x PBS, 30% glycerol, 30% ethylene glycol) and stored at −20 °C until staining.

ISH procedures were performed as previously described for ISH-HCR ^20,49^. The probes for ISH-HCR were designed to minimize off-target complementarity using a homology search by NCBI Blastn (https://blast.ncbi.nlm.nih.gov), and they were designed to have split-initiator sequences with an mRNA binding site. The number of target sites per single mRNA varied from 5 to 20 (Supplementary Table 1) based on their abundance. The DNA probes were synthesized as standard desalted oligos (Integrated DNA Technologies).

The sections were washed with PBS containing 0.2% Triton (PBST) and immersed in methanol with 3% hydrogen oxide for 10 min, followed by PBST washing for 5 min twice. After washing, the sections were prehybridized for 5 min at 37 °C in a hybridization buffer (IPL-HB, Nepagene). For multi-round ISH, the sections were treated with another hybridization solution containing a mixture of 10 nM probes for *Esr1*, *Moxd1, Penk*, *neurotensin*, and *Vglut2* mRNAs, and these probes contained the split-initiator sequences for amplification of S86, S73, S23, S72, and S45 hairpin DNAs (Supplemental Table 1), respectively. For single round ISH, hybridization buffer contained 10 nM of *Penk*, *Drd1* and *Tacr1* with the split-initiator sequences for amplification of S23, S41 and S72 hairpin DNAs. The specimens were incubated in the hybridization buffer overnight at 37 °C. After hybridization, the sections were washed three times for 10 min in 0.5× SSC containing 0.1% Tween 20 at 37 °C. Then, the sections were bleached by an LED illuminator (TiYO, Nepagene) for 60 min in PBST.

For HCR amplification, hairpin DNA solutions were separately snap-cooled (heated to 95 °C for 1 min and then gradually cooled to 65 °C for 15 min and 25 °C for 40 min) before use. The sections were incubated in amplification buffer (IPL-AB, Nepagene) for 5 min and then immersed in another amplification buffer containing 60 nM hairpin DNA and Hoechst33342 (2 μg/ml, Dojindo, H342) for 2 hours at 25 °C. After washing with PBST, the samples for single round ISH were coverslipped with mounting media containing antifade (1% n-propyl gallate and 10% Mowiol4-88 in PBS).

For multi-round ISH, the samples were coverslipped with 50% glycerol/PBS. Fluorescence images were acquired by confocal microscopy, and then, the sections were bleached with the LED illuminator for 120 min in PBST. During bleaching, the cover glass was removed from the sections. The sections were subjected to the next round of HCR reaction, and the imaging and bleaching procedures were repeated similarly. After third round HCR amplification, the sections were processed for c-Fos immunohistochemistry (IHC).

IHC was performed using an identical protocol irrespective of ISH pretreatment. Samples were washed in PBST, blocked for 30 min in 0.8% Block Ace/PBST, and then incubated overnight at 4 °C in 0.4% Block Ace/PBST containing one of the following primary antibodies or their combination: rabbit anti-c-Fos (sc-52, Santa Cruz), rabbit anti-tyrosine hydroxylase (AB152, Millipore), rabbit anti-5-HT (S5545 Sigma), chicken anti-GFP (ab13970, Abcam), and goat anti-mCherry (AB0040-200, SICGEN). After three washes in PBST, samples were incubated for 1 h in PBST containing one of the following secondary antibodies or their combination: donkey anti-rabbit Alexa Fluor 647 (ab150067, abcam), donkey anti-goat Alexa Fluor 568 (ab175704, abcam) and donkey anti-chicken Alexa Fluor 488 (703-545-155, Jackson ImmunoResearch). Sections were then washed three times in PBS and coverslipped.

Fluorescent photomicrographs were obtained using a Nikon Eclipse Ni microscope equipped with the A1R confocal detection system under 20×/0.75 NA objective lenses at 16 μs/pixel speed at 0.314 μm/pixel resolution for ISH-HCR sections (Nikon Instruments Inc., Tokyo, Japan). Tiled images were captured and automatically stitched by NIS-Elements C software (Nikon Instruments Inc., Tokyo, Japan). Images were analyzed using ImageJ software (version 1.50i, NIH, USA) to adjust the contrast. Quantification of the fluorescent photographs was performed at the identical threshold and adjustment of contrast for the same staining samples. In the multi-round ISH-IHC, the XY position of the images was adjusted manually based on the nuclear staining. Cells were considered mRNA-positive when two or more granular signals were observed adjacent to the nucleus, and their positions were recorded manually. For c-Fos immunoreactivity, cells were judged positive when signals exceeding a defined threshold were detected in the nuclear profile. The positions of the positive cells recorded using the above method were compiled based on their coordinates, and a cell map was reconstructed.

### Behavioral Assays

All behavioral assays were conducted 1–2 h after the onset of the dark phase. Male behavior was recorded under near-infrared illumination using an infrared-sensitive digital video camera (HC-VX985M, Panasonic). To handle the mice for cage introduction, dim red light was turned on for several seconds before and after introducing the mouse. Most behavioral recordings were scored in 1-s bins, whereas fiber photometry–related behavioral recordings were scored frame-by-frame (30 fps). Male sexual behaviors were recorded as sniffing, mounting, mounting with repeated pelvic thrusts (intromission), and ejaculation. Ejaculation was identified when prolonged immobility lasting several tens of seconds was observed at the end of intromission. Intermale interactions such as biting, chasing, tumbling, and wrestling were categorized as aggressive behavior without distinguishing subtypes. Behaviors that appeared interrupted but resumed within 1 s were considered a single behavioral event.

In pharmacogenetic and optogenetic experiments, sexual behavior was observed for 30 min after the introduction of a female mouse. In pharmacogenetic experiments, the introduced female was left in the cage until the following day, and the presence of a copulatory plug was checked even in males that did not display sexual behavior. In optogenetic experiments, the female was removed after the 30-min observation period. For other sexual behavior observations, males were monitored for at least 60 min after female introduction. When ejaculation occurred, the trial was classified as a full mating episode, whereas trials without ejaculation were classified as partial mating episodes. In the behavioral observations toward an anesthetized female (Figure 4l–4n), the female was lightly anesthetized by pentobarbital (20mg/kg i.p.) 10 minutes before introduction. Male behavior toward the female was recorded for 5 minutes. Male-male interactions were limited to 10 min after the introduction of a male mouse in order to prevent injuries resulting from excessive aggressive behavior. All behavioral observations were performed according to previously reported procedures ^50,51^.

### Chemogenetic manipulation

In chemogenetic experiments, mice received intraperitoneal drug administration 30 min before the start of each experiment. During two sexual behavior training sessions, PBS was intraperitoneally injected to habituate the animals to the injection procedure. After confirming that sexual behavior occurred normally in two consecutive tests, a total of four sexual behavior experiments were conducted under a schedule of weekly injections alternating between PBS and CNO. Following the sexual behavior experiments, four aggression behavior experiments were conducted under the same weekly alternating PBS and CNO injection schedule. For CNO administration, mice expressing hM3Dq received intraperitoneal injections of 0.5 mg/kg CNO, whereas mice expressing hM4Di received intraperitoneal injections of 1.25 mg/kg CNO. In sexual behavior tests, USVs of mice were recorded for 1 min immediately after the introduction of a female mouse using an ultrasonic recordable microphone (Ultramicrophone, Noldus) and Audacity software. Recorded sounds were processed to exclude the audible range (<50 kHz), and syllables of USVs were automatically detected and their number was counted using USVSEG software ^52^.

### Optogenetic manipulation

In optogenetic experiments, light stimulation was delivered using a blue LED (465–485 nm, Xlamp XQ-E, CREE) as the light source. Constant-current pulses were generated by an Arduino controller, and stimulation was applied at 20 Hz with a pulse width of 10 ms. The light intensity was adjusted to exceed 4 mW at the fiber tip. The LED was secured above the intracranially implanted optical fiber on a 3D-printed base, and the power-supply cable was connected to the power source through a slip ring. After surgery, mice were maintained with the LED and power cable connected to the head until the end of the experiment, and during the experimental sessions the cable was connected to the power supply. Behavioral experiments were conducted once per week, beginning one week after more than two normal sexual behaviors were observed during weekly sexual behavior training. In animals in which the cMPOA was stimulated, control experiments without stimulation were performed in the first and fourth weeks. In the second week, light stimulation was performed after the introduction of a female into the cage. Light stimulation was applied five times at 2-min intervals, each lasting 10 s. In the third week, light stimulation was performed before the introduction of a female into the cage. Light stimulation was applied 20 times at 30-s intervals, starting 20 min prior to female introduction, with each stimulation lasting 10 s. In mice in which axon terminals in the VTA or PAG were optically stimulated, control experiments without stimulation were performed in the first and third weeks, whereas pre-introduction stimulation was applied in the second and fourth weeks prior to female introduction. The stimulation parameters for pre-introduction were identical to those used for cMPOA stimulation.

### Data analysis

No statistical methods were used to determine the sample size. Sample sizes were selected to reliably measure experimental parameters, conform to standards in relevant fields, and minimize the number of animals used while remaining compliant with ethical guidelines. Randomization was applied to determine the order of the experiments and the group assignments. Genotype information and details of surgical procedures were fully blinded during all analyses, and histological analyses, behavioral experiments, and the quantification and analysis of video data were performed with anonymized subjects. Extraction of primary data from images or videos was conducted by an individual different from the experimenter. Statistical analyses were performed using R software, with the CAR package applied when necessary. Categorical frequency data comparing behavioral patterns between two groups of mice (Figure 1A, 6G) were analyzed using Fisher’s exact probability test. For independent continuous data comparison, analyses were performed using Welch’s t-test or Welch’s ANOVA without assuming equal variance. For paired data, comparisons were conducted using paired t-tests. Behavioral data for chemogenetics and optogenetics were analyzed by general linear model in which individual subjects were included as a factor. The significance of the coefficient estimates in the general linear model was tested using Type II-ANOVA. All statistical analyses were performed using two-sided tests. Correction of *p* values for multiple comparisons was performed using Shaffer’s method.

## Acknowledgements

We thank Hiroko Arai, Akane Iijima, Naomi Ohno, Harumi Kageyama, Emi Takeuchi for research support.

This work was supported by JSPS KAKENHI (18K06509, 21K06414, 24K09673 to Y. T.; 20H00567, 22K19410, 24H00671, 24H01219 to H.F), Research Grant from Narishige Neuroscience Research Foundation (to Y. T.), Sumitomo Foundation for Basic Science Research Projects (to Y.T.), Medical Research Grants of Takeda Science Foundation (to Y.T.), Uehara Memorial Foundation (to H. F.), Mitsubishi Foundation (to H. F.), the Chugai Foundation for Innovative Drug Discovery Science (to H. F.), the Naito Foundation (to H. F.), Takeda Science Foundation (to H. F.), and Toho University Grant for Research Initiative Program (TUGRIP)(to H. F.), and the Science Research Promotion Fund (to H. F).

## Author Contributions

YT: conceptualization, methodology and investigation.

YT, KK and KN: formal analysis. YT and HF: writing, funding acquisition, and supervision.

## Competing interests

The authors declare no competing interests.

## Materials & Correspondence

Material requests should be addressed to: Yousuke Tsuneoka, PhD.

Hiromasa Funato. MD, PhD.

## Data availability

The data in the current study are available from the corresponding author upon request.

## Supplementary figures

**Figure S1.**
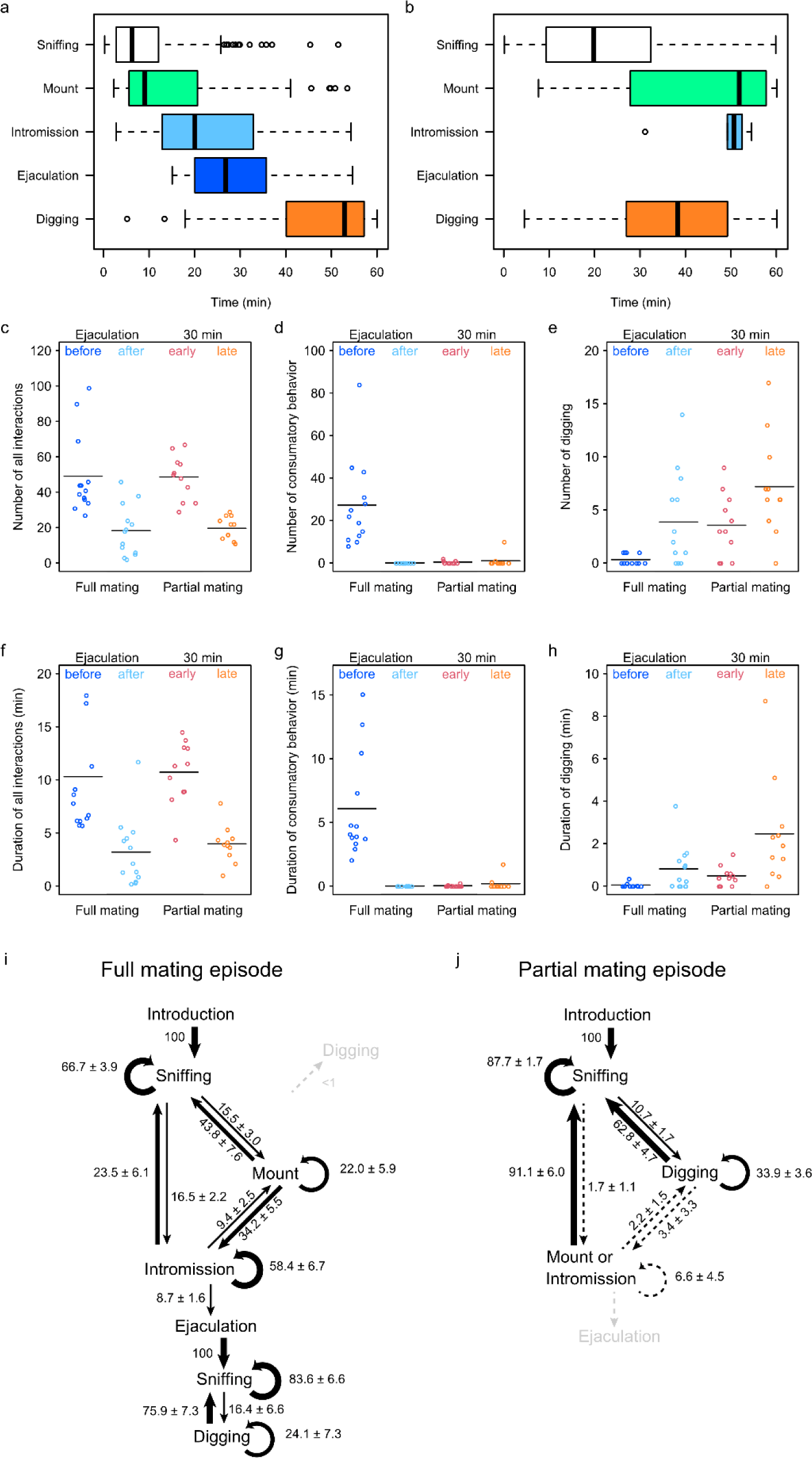
Behavioral architecture and transition dynamics of full and partial mating episodes. (a, b) Box plots showing timing of observed sexual behavior events. (a) Full mating episode (n = 13 mice). (b) Partial mating episode (n = 11 mice). (c–h) Comparison of male behaviors between full mating episodes (n = 13 mice) and partial mating episodes (n = 11 mice). Behaviors were categorized relative to ejaculation (before vs. after), and relative to the 30-min time point during a 60-min observation for the partial mating group (early vs. late phase). (c) total number of interactions. (d) number of consummatory behavior. (e) number of digging bouts. (f) duration of interactions. (g) duration of consummatory behavior. (h) duration of digging behavior. Horizontal bar indicates mean. (i,j) Transition probabilities of male behaviors following female introduction during full mating episodes (i) (n = 13 mice) and partial mating episodes (j) (n = 11 mice). Numbers indicate the probabilities of behavioral transitions (%; mean ± s.e.m.). Line thickness corresponds to the magnitude of the transition probability.

**Figure S2.**
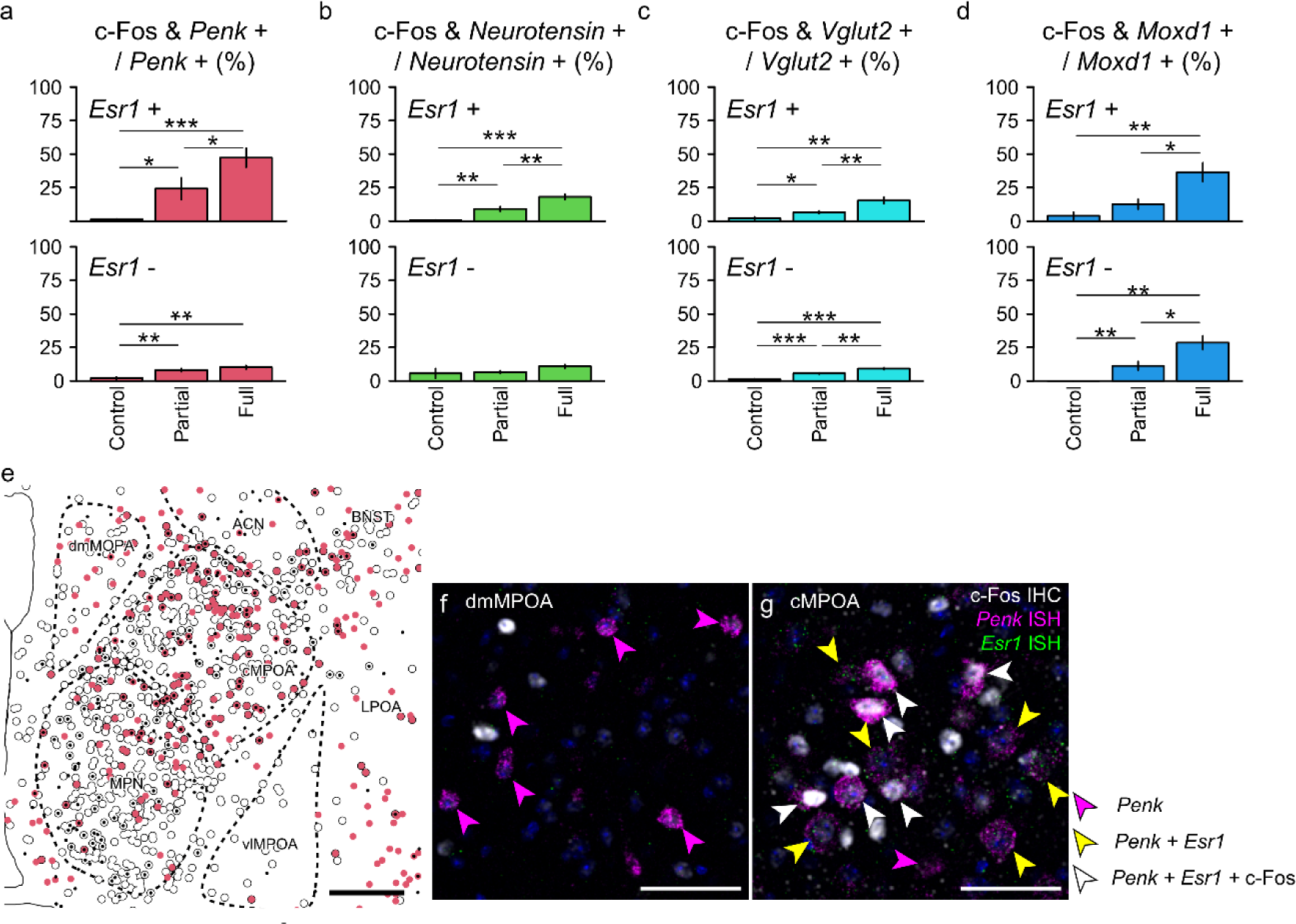
Differential activation profiles of MPOA neuronal subtypes during full and partial mating episodes. (a–d) Proportion of c-Fos–positive cells within each subtype, separated into *Esr1*-positive (top) and *Esr1*-negative (bottom) populations. (a) *Penk*-positive cells. (b) *Neurotensin*-positive cells. (c) *Vglut2-*positive cells. (d) *Moxd1*-positive cells. Data are expressed as mean ± s.e.m. (control: n = 4 mice, partial mating: n = 8 mice, full mating: n = 7 mice). Welch’s *t*-test followed by Shaffer’s correction. * *P* < 0.05, ** *P* < 0.01, *** *P* < 0.001. (e) Representative reconstruction of the distribution of c-Fos, *Penk* and *Esr1* positive neurons. Small black dots indicate c-Fos signals, red circles indicate *Penk* positive cells, and black outlines denote *Esr1*-positive cells. ACN, anterior commissural nucleus; BNST, bed nucleus of the stria terminalis; cMPOA, central part of the medial preoptic area; dmMPOA, dorsomedial part of the medial preoptic area; LPOA, lateral preoptic area; MPN, medial preoptic nucleus; vlMPOA, ventrolateral part of the medial preoptic area. (f,g) Representative merged images of c-Fos IHC with *Penk* and *Esr1* mRNA in situ hybridization (ISH). In the dorsomedial MPOA, few double- or triple-positive cells were observed (f), whereas in the cMPOA many double- and triple-positive cells were observed (g). Magenta arrowheads indicate *Penk* single positive cells, yellow arrowheads indicate *Penk/Esr1* double-positive cells, and white arrowheads indicate *Penk*/*Esr1*/c-Fos triple-positive cells. Nuclei were counterstained with Hoechst 33342. Scale bars: 50 µm.

**Figure S3.**
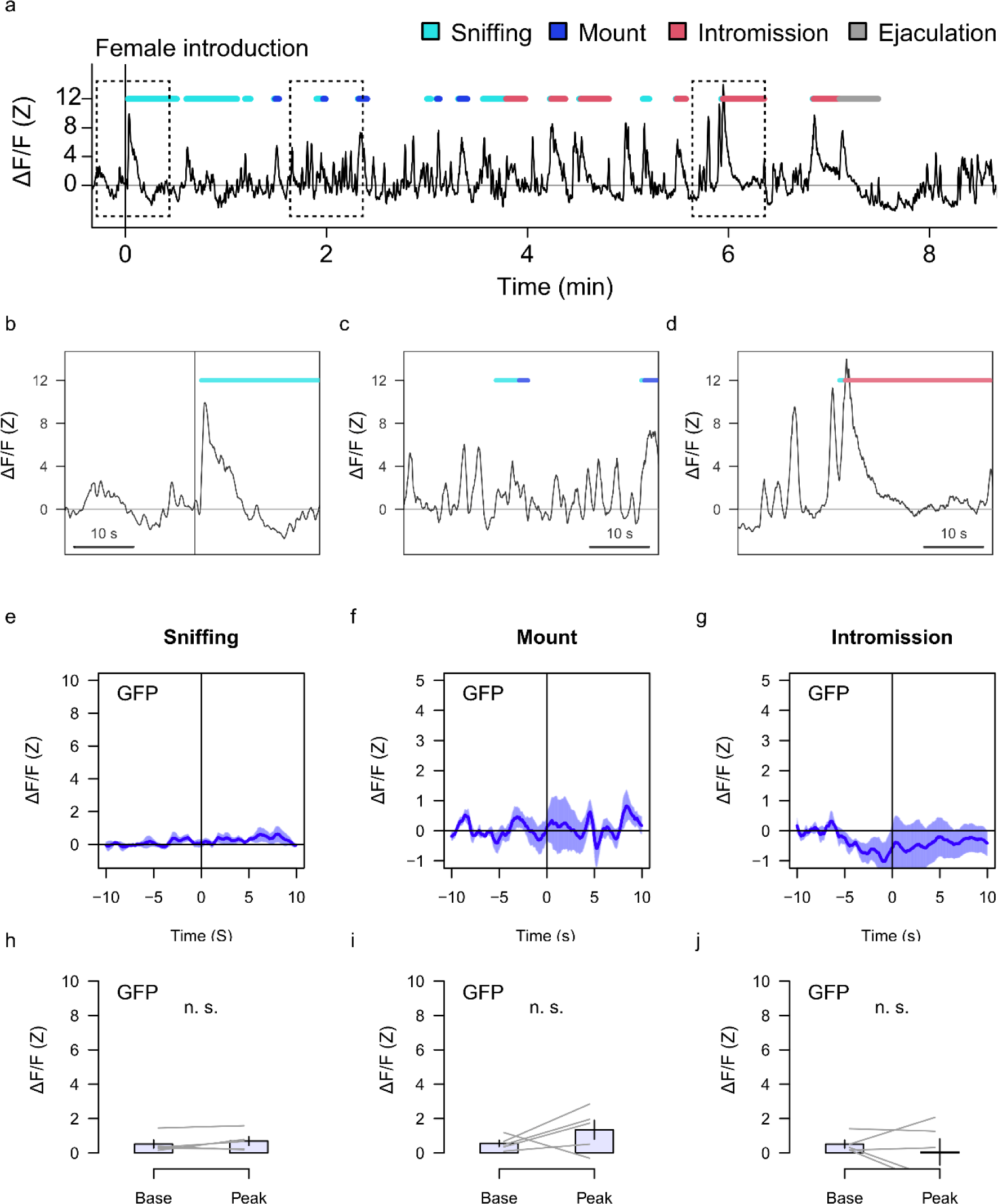
Fiber photometry control of cMPOA-Penk neurons using GFP during male sexual behavior. (a–d) Representative calcium imaging ΔF/F traces from a male mouse expressing GFP in cMPOA Penk neurons during a full mating episode. Behavioral events are indicated by color-coded bars above the traces. (a) Overview from female introduction to ejaculation. Dashed boxes indicate regions shown in higher magnification in b-d. (b) Period surrounding female introduction. (c) Period surrounding sniffing events. (d) Period surrounding intromission. (e–g) Peri-event time histogram (PETH) of ΔF/F signals aligned to the onset of sniffing (e), mount (f), an intromission (g). in control mice expressing GFP (n = 5 mice). Shaded areas indicate s.e.m. (h-j) Comparison of peak ΔF/F values between baseline and post-onset for sniffing (h), mounting (i) and intromission (j). No significant differences were observed (n = 5 mice, two-sided paired *t*-test, *P* = 0.295, 0.281 and 0.532, respectively). Data are expressed as mean ± s.e.m.

**Figure S4.**
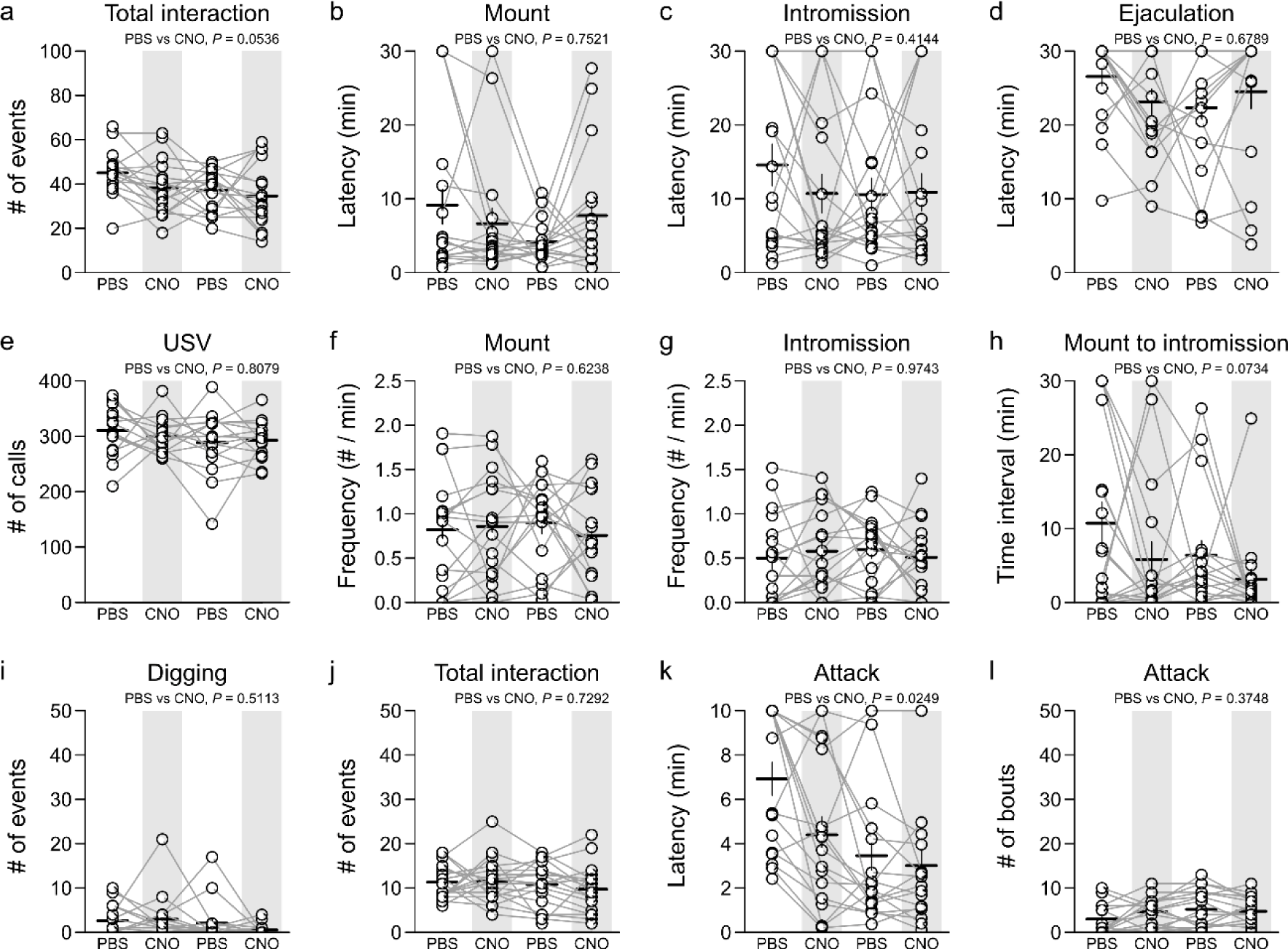
CNO administration does not alter male sexual behavior in GFP-expressing control mice. (a-i) Effects of CNO administration (0.5 mg/kg) on male sexual behavior in *Penk*-Cre mice expressing GFP in the cMPOA (n = 17). Total number of interactions (a). Latency to the first occurrence of mount (b), intromission (c) and ejaculation (d). Frequency of USV calls (e), mounts (f), and intromissions (g). Time interval from the initial mount to the initial intromission (h). Frequency of digging bouts (i). Horizontal and vertical bars indicate means and s.e.m., respectively. Group differences were tested by type II ANOVA. (j-l) Male–male interactions following CNO administration. Total number of interactions (j), latency to the first occurrence of attack (k), and number of attacks (l). Horizontal and vertical bars indicate means and s.e.m., respectively. Lines connect observations from the same mice under PBS and CNO conditions. Group differences were tested by type II ANOVA.

**Figure S5.**
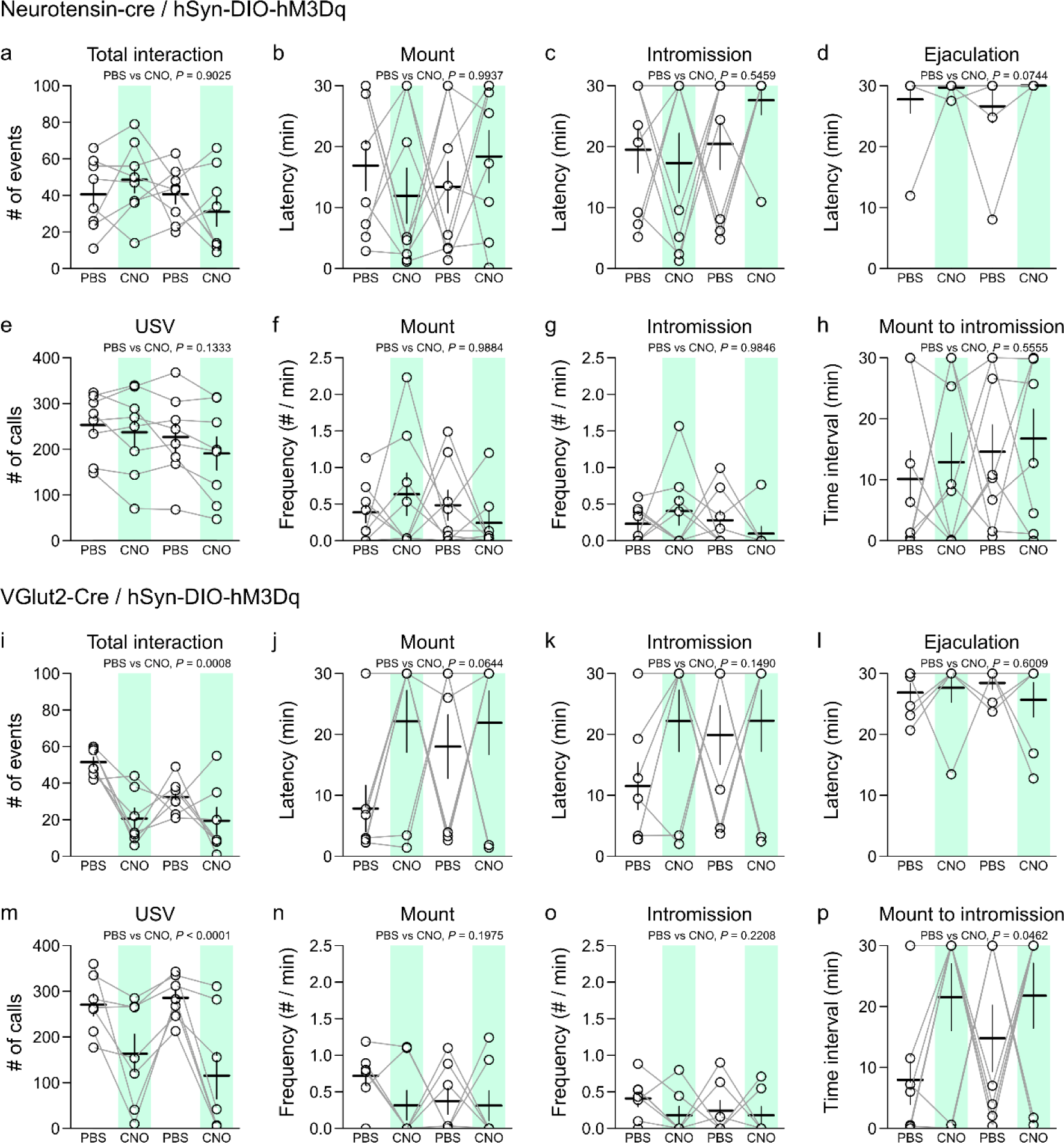
Chemogenetic activation of MPOA neurotensin+ or Vglut2+ neurons does not facilitate male sexual behavior. (a–h) Effects of CNO administration on male sexual behavior of neurotensin-Cre mice expressing hM3Dq in the MPOA (n = 7 mice). Total number of interactions (a). Latency to the first occurrence of mount (b), intromission (c) and ejaculation (d). Frequency of USV calls (e), mounts (f), and intromissions (g). Time interval from the initial mount to the initial intromission (h). (i-p) Effects of CNO administration on male sexual behavior of Vglut2-Cre mice expressing hM3Dq in the MPOA (n = 7 mice). Total number of interactions (i). Latency to the first occurrence of mount (j), intromission (k) and ejaculation (l). Frequency of USV calls (m), mounts (n), and intromissions (o). Time interval from the initial mount to the initial intromission (p). Horizontal and vertical bars indicate means and s.e.m., respectively. Lines connect observations from the same mice under PBS and CNO conditions. Group differences were tested by type II ANOVA.

**Figure S6.**
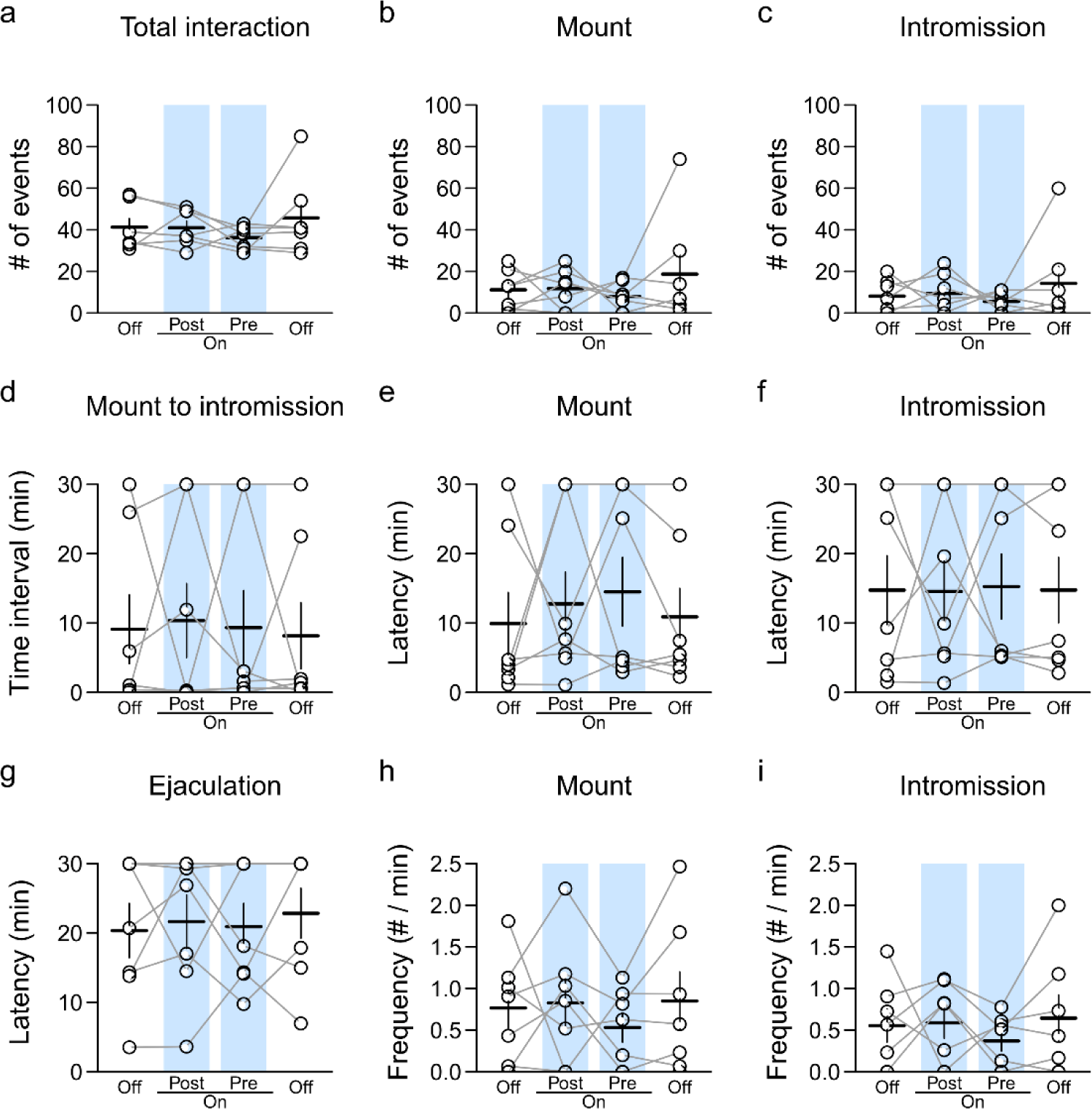
Lack of behavioral effects of light stimulation to cMPOA Penk+ neurons expressing GFP. (a–i) Effects of light stimulation on male sexual behavior in Penk-Cre mice expressing GFP in the cMPOA (n = 7 mice). Total number of interactions (a). Number of events for mount (b) and intromission (c). Time interval from the initial mount to the initial intromission (d). Latency to the first occurrence of mount (e), intromission (f) and ejaculation (g). Frequency (events per minute) of mount (h) and intromissions (i). Horizontal and vertical bars indicate means and s.e.m., respectively. Lines connect observations from the same mice under PBS and CNO conditions. Group differences were tested by type II ANOVA.

**Figure S7.**
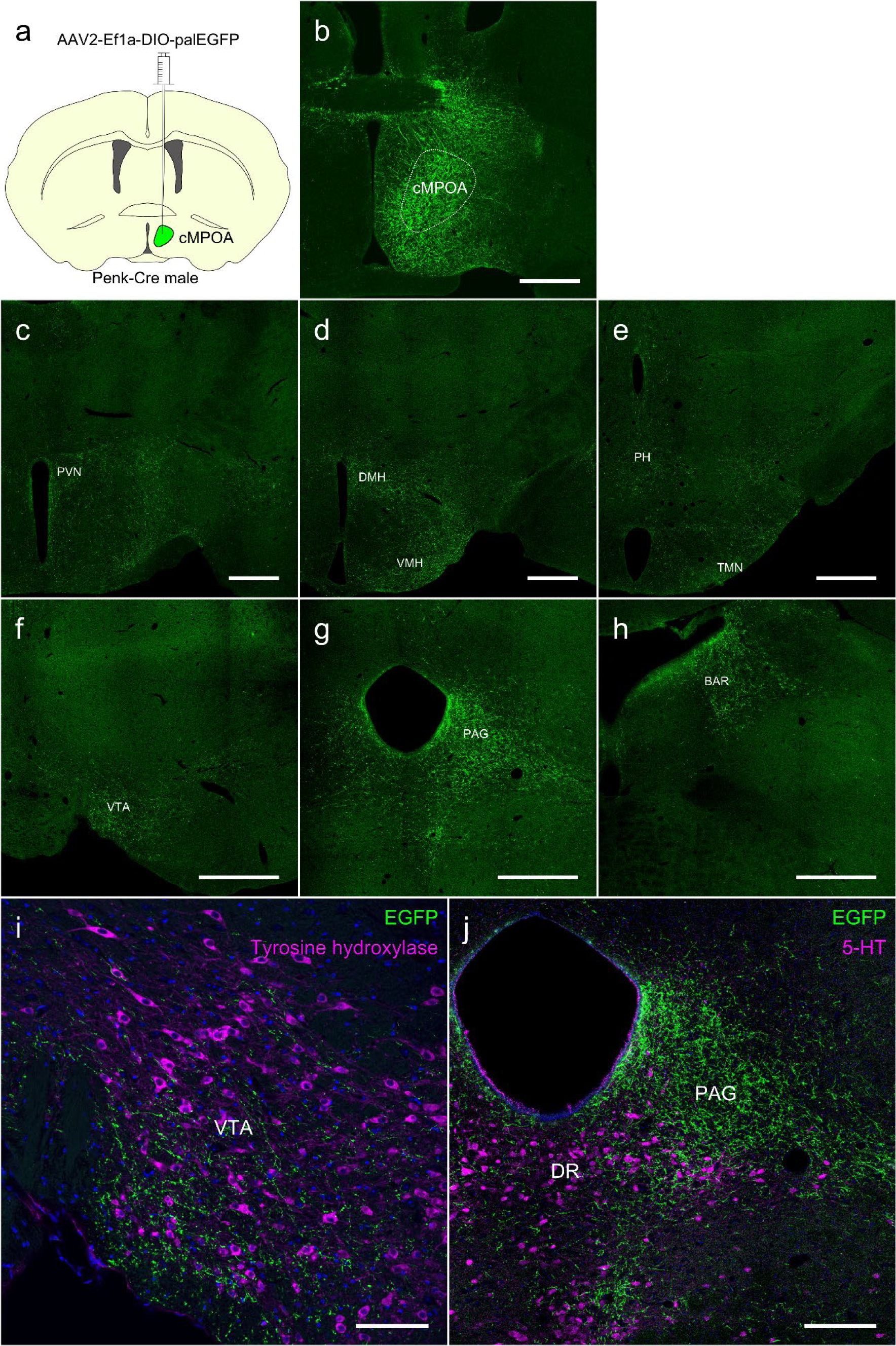
cMPOA Penk+ neurons project to multiple hypothalamic and midbrain regions. (a) Schematic illustration of Cre-dependent palmitoylated EGFP-expressing AAV injection into the cMPOA in Penk-Cre male mice. (b) Representative fluorescence photomicrograph showing the AAV injection site in the MPOA. (c-h) Representative fluorescence photomicrographs of GFP-positive fiber projection areas. GFP-positive fibers were found in the PVN (c), DMH and VMH (d), PH and TMN (e), VTA (f), PAG (g) and BAR (h).(i, j) Double immunofluorescence images of EGFP (green) and tyrosine hydroxylase (magenta) in the VTA (i) or 5-HT (magenta) in the DR (j). BAR, Barrington’s nucleus. DMH, dorsomedial hypothalamic nucleus. DR, dorsal raphe nucleus. PH, posterior hypothalamic nucleus. PVN, paraventricular nucleus. TMN, tuberomammillary nucleus. VTA, ventral tegmental area. VMH, ventromedial hypothalamic nucleus. Scale bars: 500 µm (b-h), 100 µm (i), and 200 µm (j).

## Notes

### Competing Interest Statement

The authors have declared no competing interest.

